# The ion channel TRPM8 is a direct target of the immunosuppressant rapamycin in primary sensory neurons

**DOI:** 10.1101/2024.01.24.577111

**Authors:** José Miguel Arcas, Khalid Oudaha, Alejandro González, Jorge Fernández-Trillo, Francisco Andrés Peralta, Júlia Castro-Marsal, Seyma Poyraz, Francisco Taberner, Salvador Sala, Elvira de la Peña, Ana Gomis, Félix Viana

## Abstract

**Background and Purpose:** The mechanistic target of rapamycin (mTOR) signaling pathway is a key regulator of cell growth and metabolism. Its deregulation is implicated in several diseases. The macrolide rapamycin (RAP), a specific inhibitor of mTOR, has immunosuppressive, anti-inflammatory and antiproliferative properties. Recently, we identified tacrolimus, another macrolide immunosuppressant, as a novel activator of TRPM8 ion channels, involved in cold temperature sensing, thermoregulation, tearing and cold pain. We hypothesized that RAP may also have agonist activity on TRPM8.

**Experimental approach:** Using calcium imaging and electrophysiology in transfected HEK293 cells and wildtype or *Trpm8* KO mouse DRG neurons, we characterized RAP effects on TRPM8. We also examined the effects of RAP on tearing in mice.

**Key Results:** Micromolar concentrations of RAP activate rat and mouse TRPM8 directly and potentiate cold-evoked responses. These effects were also observed in human TRPM8.

In cultured mouse DRG neurons, RAP evoked an increase in intracellular calcium almost exclusively in cold-sensitive neurons. Responses were drastically blunted in *Trpm8* KO mice or by TRPM8 antagonists. Cutaneous cold thermoreceptor endings were also activated by RAP. Topical application of RAP to the eye surface evokes tearing in mice by a TRPM8-dependent mechanism.

**Conclusion and implications:** These results identify TRPM8 cationic channels in sensory neurons as novel molecular targets of the immunosuppressant RAP. These findings may help explain some of its therapeutic effects after topical application to the skin and the eye surface. Moreover, RAP could be used as an experimental tool in the clinic to explore cold thermoreceptors.

**Bullet point summary:** WHAT IS ALREADY KNOWN

- TRPM8 is a polymodal channel involved in cold detection, thermoregulation, tearing and cold pain
- Tacrolimus, a macrolide immunosupressor, is an agonist of cold-activated TRPM8 channels

WHAT THIS STUDY ADDS

- The macrolide rapamycin also activates directly TRPM8 channels in mouse sensory neurons and human TRPM8
- Rapamycin stimulates tearing in mice in a TRPM8-dependent manner CLINICAL SIGNIFICANCE
- Rapamycin, an FDA-approved drug, shows agonist activity on TRPM8 channels
- Beneficial effects of rapamycin and other macrolides on inflammatory ocular disorders may involve TRPM8 activation

## Introduction

Rapamycin is a natural macrolide compound, isolated in 1975 from the bacterium *Streptomyces hygroscopicus* in soil samples from Rapa Nui, also known as Easter Island (Vezina et al., 1975). It was originally identified as an antifungal agent and later shown to have immunosuppressive properties (reviewed by Benjamin et al., 2011). Rapamycin (sirolimus) and analogs, called rapalogs, have important clinical applications, including their use as immunosuppressants to prevent organ rejection and as tumor suppressors. Everolimus (RAD001), a derivative of rapamycin with improved water solubility, is an FDA approved drug for the treatment of various solid tumors, including neuroendocrine tumors, breast cancer, renal cell carcinoma and subependymal giant cell astrocytoma (reviewed by Li et al., 2014; Seto, 2012). Topical application of rapamycin is beneficial in ocular inflammatory disorders, including dry eye disease (Spatola et al., 2018). In addition, rapamycin has been used extensively as a research tool (reviewed by Kunz & Hall, 1993).

Rapamycin acts by inhibiting the mechanistic target of rapamycin (mTOR) (Heitman et al., 1991; Johnson et al., 2013), a key kinase for cellular growth and proliferation, sensing and integrating multiple metabolic signals: growth factors, nutrients, metabolites, and oxygen levels (Kennedy & Lamming, 2016). Rapamycin, and derivatives, bind to its receptor, FKBP12, which directly interacts with the mTORC1 complex, inhibiting its downstream signalling. The inhibition of mTORC1 disrupts glycolytic pathways, decreases lipolysis and induces autophagy, inhibiting cell proliferation (Saxton & Sabatini, 2017). Pharmacological inhibition of mTORC1 is also being explored as neuroprotectant in various neurodegenerative diseases.

Rapamycin administration has multiple effects on organism biology, which are still poorly understood and require further study. The recent discovery that rapamycin can extend the lifespan in numerous species, from yeast to mice, and improve age-related functional decline in them, has generated great excitement in the field of aging biology (Bitto et al., 2016; Harrison et al., 2009; Wilkinson et al., 2012). Indeed, this is one of the few examples of a clinically approved drug capable of slowing the aging process in mammals.

Recently, we identified tacrolimus (FK-506), a different macrolide immunosuppressant targeting calcineurin, as an agonist of TRPM8 (Arcas et al., 2019), a cold- and menthol-activated cationic channel (McKemy et al., 2002; Peier et al., 2002) expressed in the soma of a subset of small diameter primary sensory neurons and their peripheral terminals (Dhaka et al., 2008; Takashima et al., 2007). In mammals, TRPM8 plays a major role in cold temperature detection, thermoregulation and cold pain (reviewed by Almaraz et al., 2014). Moreover, TRPM8 agonists promote ocular tearing and blinking (Parra et al., 2010; Quallo et al., 2015), and can ameliorate the symptoms of dry eye disease. In addition, TRPM8 is overexpressed in prostate cancers and other solid tumors, making it an interesting target and prognostic marker for the disease (Bidaux et al., 2016; Zhang & Barritt, 2006).

Here, we asked whether rapamycin and its analog, everolimus, share agonist activity on TRPM8. We found that both activate recombinant TRPM8 channels from various species, including humans, and mouse TRPM8-expressing sensory neurons, generalizing macrolide immunosuppressants as a novel, clinically relevant, class of mammalian TRPM8 modulators.

## Methods

### Animals

Studies were performed on young adult (1 to 4 months old) mice of either sex. Mice were bred at the Universidad Miguel Hernández Animal Research Facilities (ES-119-002001) and kept in a barrier facility under 12/12 h light dark cycle with food and water *ad libitum*. Wild type (WT) mice were of the C56Bl/6J strain. All experimental procedures were performed according to the Spanish Royal Decree 53/2013 and the European Community Council directive 2016/63/EU, regulating the use of animals in research.

Two transgenic mouse lines were used for calcium imaging experiments and electrophysiological recordings on DRG cultures. In *Trpm8^BAC-EYFP^* mice, the fluorescent protein EYFP is expressed under the *Trpm8* promoter (Morenilla-Palao et al. 2014). For experiments with *Trpm8* KO mice, we used a transgenic knockin line, *Trpm8^EGFPf^*, in which the *Trpm8* locus was disrupted and farnesylated EGFP was inserted in frame with the *Trpm8* start codon (Dhaka et al. 2007). Homozygous mice *(Trpm8^EGFPf/EGFPf^*) are null for TRPM8 (Dhaka et al., 2007; Parra et al., 2010). As previously described, to enhance EGFPf expression the lox-P-flanked neomycin selection cassette introduced into the *Trpm8* locus during the generation of the transgene was excised (Dhaka et al., 2008). Both transgenic lines allowed the identification of TRPM8-expressing cells by the expression of EYFP or EGFP fluorescence. Moreover, *Trpm8^EGFPf/+^* (i.e. heterozygous) allowed recordings from EGFP(+) neurons with one functional copy of TRPM8. The genotype of transgenic mice was established by PCR.

Experiments in cultured DRG neurons and behavioral experiments were performed on animals of either sex. Experiments on the firing of mouse afferents (i.e. skin-nerve preparation) were performed on male mice only due to limited availability of female mice at the time. TRPM8 channels are expressed in both sexes and the effect of rapamycin on TRPM8 channels is direct. We did not observe any differences in the results between male and female animals, so the data were pooled.

### Culture and transfection of mammalian cell lines

Human embryonic kidney 293 cells (HEK293) were maintained in DMEM plus Glutamax, supplemented with 10% fetal bovine serum, 100 U/ml penicillin (Gibco) and 100 μg/ml streptomycin (Gibco), incubated at 37 °C in a 5% CO_2_ atmosphere. HEK293 cells were plated in 24-well dishes at 2×10^5^ cells/well, and transiently transfected with Lipofectamine 2000 (Thermo Fisher Scientific). When necessary, we cotransfected the cells with 1 μg of TRPM8 channel plasmid (from different species) and 0.5 μg of EGFP or mCherry plasmid. For the transfection, 2 μl of Lipofectamine 2000 was mixed with the DNA in 100 μl of OptiMem (Thermo Fisher Scientific), a reduced serum media. Electrophysiological and calcium imaging recordings took place 24-36 hours after transfection. The evening before the experiment, cells were trypsinized (0.25% trypsin-EGTA) and reseeded at lower density in 12 mm diameter glass coverslips previously treated with poly-L lysine (0.01%, Sigma-Aldrich).

The expression vectors used and their source were as follows: mouse TRPM8 (NM_134252) in pcDNA5, kindly provided by Ardem Patapoutian (Scripps Research Institute, La Jolla, USA), was used as a wildtype TRPM8. Human TRPM8 in pcDNA3 (Veit Flockerzi, Saarland University), human TRPA1 in pCMV6-AC-GFP (Veit Flockerzi, Saarland University), rat TRPV1 in pcDNA3 (David Julius, UCSF) and human TRPV1 in pcDNA3(+) (Andreas Leffler, Hannover Medical School) were also transiently transfect in HEK293 cells using the same techniques. The menthol-insensitive mouse TRPM8-Y745H mutant was generated by site-directed mutagenesis (Malkia et al., 2009). Enhanced Green fluorescent protein (GFP) was expressed from the pGREEN LANTERN™-1 vector (Life Technologies). The mCherry plasmid was obtained from Addgene (#84329) (van Unen et al., 2016).

HEK293 cells stably expressing rat TRPM8 channels (CR#1 cells) (Brauchi et al., 2004) were cultured in DMEM containing 10% of fetal bovine serum, 100U/ml penicillin, 100 mg/ml streptomycin (Gibco) and 450 µg/ml geneticin (G418).

### DRG cultures

Adult mice of either sex (1-4 months) were anesthetized with isofluorane and decapitated. The spinal cord was isolated and dorsal root ganglia (DRG) were dissected out from all spinal segments and maintained in ice-cold HBSS solution. After isolation, DRGs were incubated with collagenase type XI (Sigma) and dispase II for 30-45 min in Ca^2+^- and Mg^2+^-free HBSS medium at 37 °C in 5% CO_2_. Thereafter, DRG ganglia were mechanically dissociated by passing 15-20 times through a 1 ml pipette tip and filtered through a 70 µm nylon filter. Neurons were harvested by centrifugation at 1200 rpm during 5 minutes. The resultant pellet was resuspended in Minimun Essential Medium (MEM) supplemented with 10 % FBS, 1% MEM-vit, 100U/ml penicillin (Gibco), 100 mg/ml streptomycin (Gibco) and plated on poly-L-lysine-coated glass coverslips. Electrophysiological and calcium-imaging recordings were performed after 12-36 hours in culture.

### Fluorescence Ca^2+^ imaging

Ratiometric calcium imaging experiments were conducted with the fluorescent indicator Fura-2 (Thermo Fisher Scientific). DRG neurons or HEK293 cells were incubated with 5 μM Fura-2 AM and 0.2% Pluronic (Thermo Fisher Scientific) (prepared from a 200 mg/ml stock solution in DMSO) for 45 min at 37 °C in standard extracellular solution. Fluorescence measurements were obtained on a Leica inverted microscope fitted with an Imago-QE Sensicam camera (PCO). Fura-2 was excited at 340 and 380 nm (excitation time 60 ms) with a rapid switching monochromator (TILL Photonics) or an LED-based system (Lambda OBC, Sutter Instruments). Mean fluorescence intensity ratios (F340/F380) were displayed online with TillVision (TILL Photonics) or Metafluor (Molecular Devices, San Jose, CA, USA) software. The standard bath solution (290 mOsm/kg) contained (in mM): 140 NaCl, 3 KCl, 2.4 CaCl_2_, 1.3 MgCl_2_, 10 HEPES, and 10 glucose, adjusted to a pH of 7.4 with NaOH. Calcium imaging recordings were performed at a basal temperature of 33 ± 1 °C. Before the start of the experiment, an image of the microscopic field was obtained with transmitted light and under 460 nm excitation wavelength, in order to identify fluorescent cells.

Responses to agonists were calculated by measuring the peak ratio values, after subtracting the mean baseline fluorescence ratio during the 15 seconds previous to agonist application. Responses were scored as positive if the increase in fluorescence (ΔFura2 ratio) was > 0.08.

### Electrophysiology in cultured cells

Whole-cell voltage- and current-clamp recordings were obtained from mouse DRG neurons or transiently transfected HEK293 cells with borosilicate glass patch-pipettes (Sutter Instruments, 4-8 MΩ resistance) and were performed simultaneously with temperature recordings. Signals were recorded with an Axopatch 200B patch-clamp amplifier (Molecular Devices) and digitized through a Digidata 1322A (Molecular Devices). Stimulus delivery and data acquisition were performed using pCLAMP9 software (Molecular Devices).

For neuronal recordings, we used the standard bath solution (see above) at a basal temperature of 33 °C. The intracellular solution (280 mOsm/kg) contained (in mM): 115 K-gluconate, 25 KCl, 9 NaCl, 10 HEPES, 0.2 EGTA, 1 MgCl, 1 Na_2_GTP and 3 K_2_ATP, adjusted to pH 7.35 with KOH. In voltage-clamp recordings amplifier gain was set at x1, sampling rate was set to 10 kHz and the signal was filtered at 2 kHz. Neurons were voltage-clamped at a potential of −60 mV. For current clamp recordings gain was set at x10, acquisition rate was 50 kHz and the signal was filtered at 10 kHz. Once in the whole-cell configuration, resting membrane potential was measured. In neurons that fired action potentials at rest, a small DC current was injected to bring the cell to around −55 mV.

For electrophysiological experiments in HEK293 cells, to minimize desensitization of TRPM8 responses, a calcium-free extracellular solution was used (in mM): 144.8 NaCl, 3 KCl, 1 EGTA, 1.3 MgCl2, 10 HEPES, and 10 glucose (290 mOsm/kg, pH adjusted to 7.4 with NaOH). The intracellular solution for HEK293 recordings was (in mM): 135 CsCl, 2 MgCl2, 10 HEPES, 1 EGTA, 5 Na_2_ATP and 0.1 NaGTP, adjusted to pH 7.4 with CsOH (280 mOsm/kg). For experiments examining the dose-dependence of rapamycin, the extracellular solutions contained (in mM): 150 NaCl, 10 EDTA, 10, HEPES, 10 glucose (332 mOsm/kg, pH adjusted to 7.4 with NaOH). In these experiments, the intracellular solution was (in mM): 150 NaCl, 3 MgCl2, 10 HEPES, 5 EGTA (300 mOsm/kg, pH adjusted to 7.22 with NaOH). Due to the pH adjustment of the EDTA stock solution with NaOH, we calculated an excess of 19 mM Na in the extracellular solution compared to the intracellular. Recordings were performed at a basal temperature of 33 ± 1°C.

After a Gigaohm seal was formed and the whole-cell configuration was established, cells were voltage-clamped at a potential of −60 mV. To estimate shifts in the voltage dependence of TRPM8 activation during cold and agonist application, current-voltage (I-V) relationships obtained from repetitive (0.33 Hz) voltage ramps (−100 to +150 mV, 400 ms duration) were fitted with Origin BotlzIV function (OriginPro version 2020. OriginLab Corporation, Northampton, MA, USA) a function that combines a linear conductance multiplied by a Boltzmann activation term:

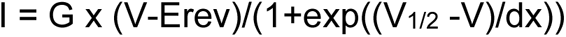

where G is the whole-cell conductance, Erev is the reversal potential of the current, V_1/2_ is the potential for half-maximal activation and dx is the slope factor.

We determined the Erev parameter independently for each condition by measuring the average voltage at which the current reversed sign, which was then kept constant during the fitting process. While the G value was shared between all conditions, V_1/2_ and dx were left free to vary for each condition. The ramps were analyzed with WinASCD software package (Prof. G. Droogmans, Laboratory of Physiology, KU Leuven).

For the experiments examining the opening and inactivation kinetics, 100 ms duration voltage steps were applied (from −80 to +240 mV) from a holding potential of −80 mV. The currents were acquired at 10 kHz, filtered at 2 kHz.

### Single channel recordings

Gating of TRPM8 channels by agonists requires the intracellular presence of PIP2 and polyphosphates (Zakharian et al., 2009). To minimize fluctuations in the concentration of these intracellular mediators, we characterized channel activity during cell-attached recordings.

Single channel recordings were obtained, at room temperature (25.2 ± 0.5 °C °C), from HEK2923 cells transiently transfected with mTRPM8 and mCherry using a HEKA EPC10 amplifier with PatchMaster software (HEKA Elektronik, Lambrecht, Germany). Borosilicate glass patch pipettes were fire-polished to a resistance of 7-10 MΩ. Current signal was sampled at 50 kHz and filtered at 3 kHz.

After obtaining a Gigaohm seal, the standard bath solution (in mM): 140 NaCl, 3 KCl, 1.2 MgCl_2_, 1 EGTA, 10 Glucose, 10 HEPES, pH: 7.4, was switched to a calcium-free solution containing 145 NaCl, 3 KCl, 10 Glucose, 10 HEPES, 1 EGTA, pH: 7.4, to minimize TRPM8 desensitization. The pipette solution contained (in mM): 140 KCl, 0.6 MgCl_2_, 10 HEPES, 1 EGTA, pH: 7.4. All-point histograms of the channel current were fitted to a double Gauss function (solid line in red). The single-channel current was calculated from the difference in the two fitted peak values. Single-channel conductance events, all points’ histograms, open probability, and other parameters were identified and analyzed using Nest-o-Patch 2.1.9.8 (https://sourceforge.net/p/nestopatch/wiki/Home/) and Clampfit 9.2 (Molecular Devices).

### Temperature stimulation

Glass coverslip pieces with cultured cells were placed in a microchamber and continuously perfused with solutions warmed at 33 ± 1 °C. The temperature was adjusted with a Peltier device (CoolSolutions, Cork, Ireland) placed at the inlet of the chamber, and controlled by a feedback device (Reid et al., 2001). Cold sensitivity was investigated with a temperature drop to approximately 18 °C. The bath temperature was monitored with an IT-18 T-thermocouple connected to a Physitemp BAT-12 micro-probe thermometer (Physitemp Instruments) and digitized with an Axon Digidata 1322A AD converter running Clampex 9 software (Molecular Devices).

### Isolated skin nerve preparation

Extracellular recordings from single cutaneous primary afferent axons in an isolated mouse skin–saphenous nerve preparation were obtained following previously published procedures (Roza et al., 2006; Zimmermann et al., 2009). In brief, adult male C57Bl/6J mice were killed by cervical dislocation and the hairy skin from either hind paw, with the saphenous nerve attached, was dissected free from underlying muscles and placed in a custom made Teflon recording chamber with the corium side up (Zimmermann et al., 2009).

The chamber containing the preparation was continuously superfused at a rate of 4 ml/min with oxygenated external solution consisting of (in mM): 107.8 NaCl, 26.2 NaHCO_3_, 9.64 sodium gluconate, 7.6 sucrose, 5.55 glucose, 3.5 KCl, 1.67 NaH_2_PO_4_, 1.53 CaCl_2_ and 0.69 MgSO_4_, which was adjusted to pH 7.4 by continuously gassing with 95% oxygen–5% CO_2_. Temperature of the solution was maintained around 34 °C with a SC-20 in-line heater/cooler system, driven by a CL-100 bipolar temperature controller (Warner Instruments).

After pulling back the perineurum with Dumont #5SF forceps, a small bundle of fibers was aspirated into a patch-pipette connected to a high gain AC differential amplifier (model DAM 80; World Precision Instruments). A reference electrode was positioned inside the chamber. Input signals were amplified, digitized (CED Micro1401-3; Cambridge Electronic Design) at 25 kHz and stored in the hard drive of a PC for off-line analysis. For recording and off-line analysis, the Spike 2 software package was used (Cambridge Electronic Design).

A small, cone-shaped piece of frozen external solution was moved slowly over the corium side of the skin and used to identify cold spots: brisk thermoreceptor fiber activity was evoked by the ice cone when in the immediate vicinity of the receptive field, and this activity stopped shortly after removing the stimulus. Cold spots identified in this way where then isolated from the surrounding tissue with a small ABS thermoplastic ring, and delivery of the subsequent cold and chemical stimuli were restricted to a circular area (5 mm diameter) of the skin. Cold stimuli were performed with solutions flowing through a Peltier system custom-designed to deliver a small volume of solution inside the ring isolating the skin area. Starting from a baseline temperature of 34-35 °C, the temperature reached ∼ 12 °C in about 50 seconds.

In control conditions, at the baseline temperature of 34-35 °C, cold thermoreceptors were silent, firing action potentials during the cooling ramp. The cold threshold was defined as the temperature corresponding to the first spike during a cooling ramp. When chemical sensitization led to the appearance of ongoing activity already at basal temperature, cold threshold was taken as the mean temperature during the 60 s preceding the start of the cooling ramp. Chemical sensitivity of single fibers was tested with consecutive applications of RAP (30 µM), followed by menthol (50 µM) after a period of wash. Chemical sensitivity during recordings at 34 °C was defined as the presence of at least 20 spikes during a period of 2 minutes before the cooling ramp. In order to explore their effects on cold sensitivity, a cold temperature ramp was also applied in the presence of RAP or menthol.

### Mathematical modelling of cold thermoreceptor activity

To simulate the effect of temperature changes and agonist applications on cold thermoreceptor activity, we used the model of cold-sensitive neurons with TRPM8 channels originally described by Patricio Orio and colleagues for cold sensitive nerve endings (Olivares et al., 2015) and later for trigeminal neurons (Rivera et al., 2021). The equations and parameters are as described in Rivera et al., except for the reversal potential of the depolarizing currents that was set to +70 mV, and that no white noise current was used in order to get more reproducible results. All 52 parameter sets described by Rivera et al. were used to simulate neurons with different temperature thresholds. The model was implemented in NEURON and controlled with Python 3.8 (Carnevale et al., 2006; Hines et al., 2009)

To emulate our experimental protocol, a double temperature ramp was applied with 120 s interval between the two; each ramp consisted in cooling from 34 to 18 °C in 12 s, staying 2 s at 18 °C and then warming back to 34 °C in 14 s. The first ramp starts at t = 30 s.

We simulated the application of a pulse of TRPM8 agonist by changing g_M8_, V_half_ or both. The agonist pulse was applied at t = 120 s and maintained for 90 s approximately. Thus, the second temperature ramp is applied while the agonist pulse is on. The total simulation time is 270 s.

In *Current-Clamp* simulations, a temperature threshold is defined as the temperature at which the first action potential is fired. The time at which an action potential is fired is given by the time it crosses the 0 mV from below, and the instantaneous firing frequency is defined as the inverse of the time between two consecutive action potentials. Thus, two temperature thresholds are measured, one for each ramp (i.e. the first in control conditions and the second in the presence of agonist). Also, two corresponding values for the maximum, and average firing frequency are measured, for the first and second temperature ramps and at the beginning of the agonist pulse.

In *Voltage-Clamp* simulations, the membrane potential is clamped at −60 mV and the *I_M_*_8_ current is recorded using the same temperature ramp and agonist pulse as described above. The peak *I_M8_* current is measured when elicited by a cold ramp, an agonist pulse, or both.

We analyzed the results of the simulations with R (R Core Team, 2022), and the Python packages Numpy (Harris et al., 2020), Scipy (Virtanen et al., 2020) and Matplotlib (Hunter, 2007).

### Behavioral assessment of tearing

Adult male and female mice, C57BL/6J (n = 15) or *Trpm8* KO (n = 15) (Dhaka et al., 2008), were anaesthetized by intraperitoneal injection of a mixture of ketamine hydrochloride (80 mg/kg, Imalgene 1000; Laboratorios Merial S.A., Barcelona, Spain) and Xylazine hydrochloride (5 mg/kg, Rompun; Bayer Hispania S.L., Barcelona, Spain). Tear flow was measured in both eyes, using phenol red threads (Zone-Quick, Menicon Pharma S.A., Illkirch Graffenstaden, France), after applications of a drop (2 µl) of 1% RAP in one eye or vehicle (8% ethanol, 2% Cremophor in saline) in the other using a graduated micropipette (Gilson Pipetman P2). Each solution was applied for 2 minutes. Thereafter, excess fluid was removed using a sterile absorbent swap (Sugi® Eyespear pointed tip, Kettencach GmbH & Co., Germany). After a rest period of 5 minutes, a phenol red thread was gently placed between the lower lid and the bulbar conjunctiva at the nasal angle during 1 min. To quantify the staining of the threads, the wetted length was measured under a stereomicroscope. The eye receiving RAP was random for each mouse. The experimenters were blind to the animal’s genetic background and treatment. After measurements, animals were euthanized according to the approved protocol.

### Chemicals

Rapamycin (RAP) and 40-*O*-(2-hydroxyethyl)-rapamycin) (everolimus) were purchased from LC laboratories (Woburn, MA) and prepared in a DMSO stock (50 mM). The required volume for experiments was dissolved in pre-warmed (50 ^°^C) control solution. When RAP, or everolimus, were added to the external solution a white cloud of precipitation appeared and gentle shaking was applied until total dissolution was obtained. Due to its poor solubility in water, a solution of 30 µM RAP was the highest concentration tested. In separate experiments, application of vehicle (0.06 %DMSO) did not activate TRPM8. Menthol (Scharlau, Spain), 1R,2S,5R)-N-(4-methoxyphenyl)-5-methyl-2-propan-2-ylcyclohexane-1-carboxamide (WS-12) was purchased from Tocris Bioscience and prepared in a DMSO stock (20 mM). AMTB *(N-(*3-Aminopropyl)-2-[(3-methylphenyl)methoxy]-*N*-(2-thienylmethyl)benzamide hydrochloride; Tocris), M8-B (*N*-(2-Aminoethyl)-*N*-[[3-methoxy-4-(phenylmethoxy)phenyl]methyl]-2-thiophenecarboxamide hydrochloride), AITC (Allyl isocyanate; Sigma), Capsaicin (8-Methyl-N-vanillyl-*trans*-6-nonenamide; Sigma) were prepared as stocks and stored at −20 °C.

### Blinding and randomization

For most *in vitro* protocols, blinding was not feasible as experiments were conducted by an individual experimenter. During *in vivo* experiments, the experimenter was blind to animal phenotype and treatment.

### Experimental design and statistical analysis

The manuscript complies with BJP’s recommendations and requirements on experimental design and analysis. All experimental groups had a size of 5 or larger.

Coverslips with plated cells or neurons were only used once (i.e. a single imaging field or a single neuron). Data were obtained from multiple experiments or animals, as described in figure legends, generally on different dates.

To estimate shifts in the voltage dependence of TRPM8 activation, current-voltage (I-V) relationships obtained from repetitive (0.33 Hz) voltage ramps (−100 to +150 mV, 400 ms duration) were fitted with a function that combines a linear conductance multiplied by a Boltzmann activation term:

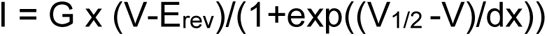

where G is the whole-cell conductance, E_rev_ is the reversal potential of the current, V_1/2_ is the potential for half-maximal activation and dx is the slope factor. The G value obtained for a high menthol concentration (800 µM) was taken as Gmax and was used for the representation of G/Gmax curves.

For the fitting of G/Gmax curves extracted from the voltage pulses protocol a Boltzmann equation was used,

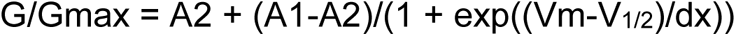

where A2 is de maximal normalized conductance, A1 is the minimal normalized conductance, Vm is the test potential, V_1/2_ is the potential for half-maximal activation and dx is the slope factor

Conductance–voltage (G–V) curves were constructed from the I–V curves of individual cells by dividing the evoked current by the driving force, according to the following equation:

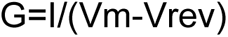

where Vm is the testing potential and Vrev is the reversal potential of the current.

The threshold temperatures were estimated as the first point at which the measured signal (F340/F380 or current) deviated by at least four times the SD of its baseline. All fittings were carried out with the Levenberg-Marquardt method implemented in the Origin 8.0 software.

Data are reported as mean ± standard error of the mean (SEM). When comparing two means, statistical significance (p<0.05) was assessed by Student’s two-tailed t test. For multiple comparisons of means, one-way ANOVAs were performed, followed by Bonferroni’s post-hoc analysis, using GraphPad Prism version 8.00 for Windows (GraphPad Software). During statistical analysis, normality of the data distribution was determined with the Shapiro-Wilk test. The decision to use parametric or non-parametric statistical tests considered additional factors. Many hypothesis tests, including ANOVA, are robust to violations of the normality assumption if the sample size is large enough. This is the case for most calcium imaging data that have samples >30. In addition, evaluation of the quantile-quantile (Q-Q) plot provided additional insight on the distribution. For nonparametric analysis we used the Kruskal-Wallis test. In the figures, statistical comparisons were marked by asterisks or an alternative symbol: * for p <0.05, ** for p < 0.01 and *** for p < 0.001.

## Results

### Rapamycin activates recombinant TRPM8 channels

To investigate the effect of RAP on TRPM8 channels, we performed intracellular Ca^2+^ imaging experiments on HEK293 cells stably expressing rat TRPM8 channels (CR#1 cells) (Brauchi et al., 2004). RAP was applied in independent experiments at four different concentrations (1, 3, 10 and 30 µM) for 150 s. As shown in Figure 1A and 1B, RAP produced a dose-dependent activation of TRPM8 with and EC_50_ of 6.5 ± 1.5 µM. RAP had poor solubility in aqueous solutions, limiting the maximum concentration that could be tested. At low concentrations the calcium response evoked by RAP had a slow rise and was sustained, while at higher concentrations RAP evoked faster responses that desensitized partially during the time of application (Fig. 1A).

**Figure 1.**
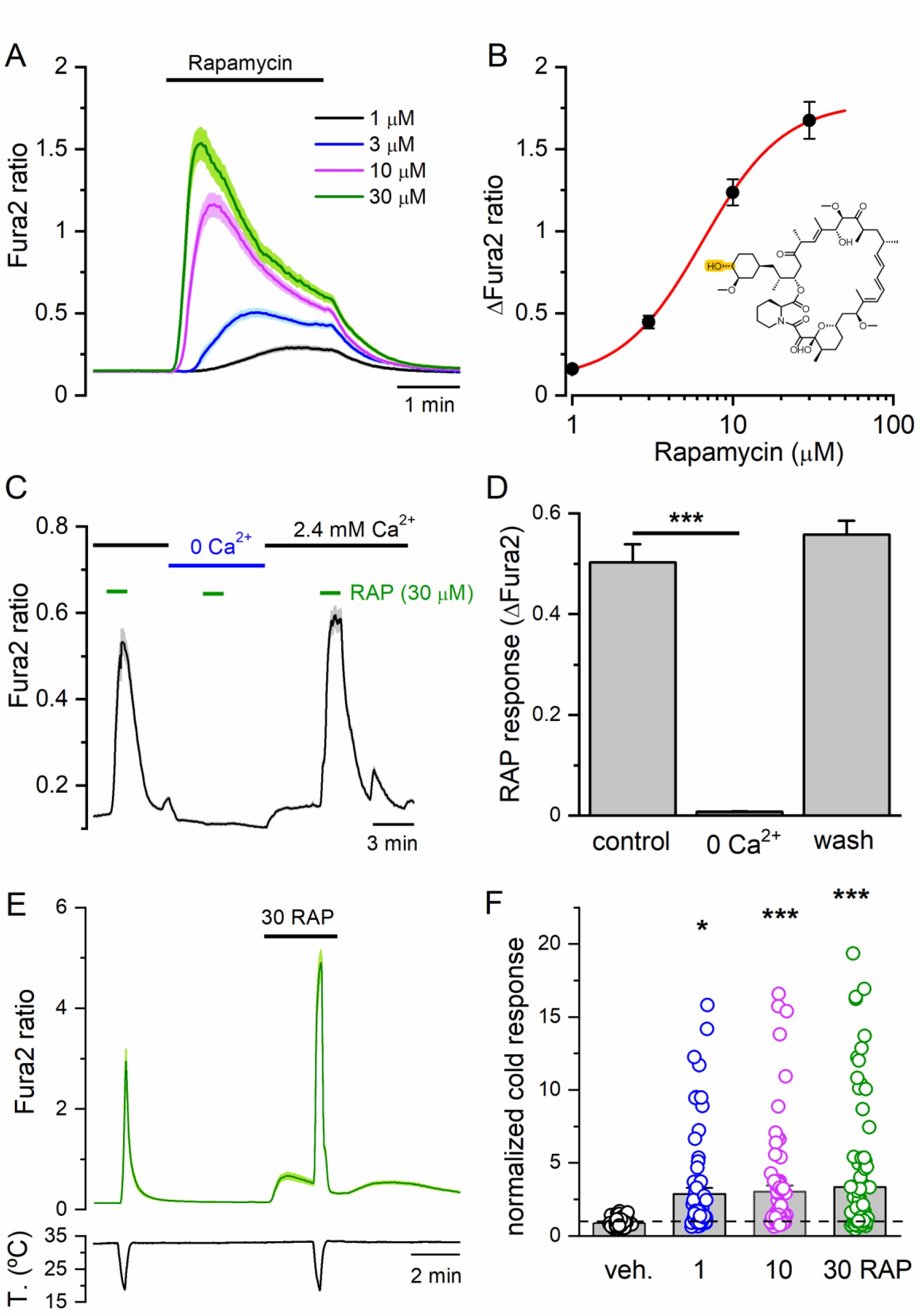
Rapamycin activates recombinant TRPM8 channels and potentiates cold-evoked responses. ***A,*** Average ± SEM Fura2 ratio time course in HEK293 cells stably expressing rat TRPM8 (CR#1 cells) during RAP application at different concentrations. A single dose was applied in individual experiments (n = 38 to 185 cells, 8 experiments). ***B,*** Dose-response curve of RAP effects. Averaged ± SEM values were obtained from individual peak amplitudes from experiments shown in A. ***C,*** Average ± SEM Fura2 ratio time course in HEK293 cells stably expressing rat TRPM8 during a protocol in which rapamycin responses were monitored in the presence (2.4 mM Ca^2+^) and in the absence (0 Ca^2+^) of extracellular calcium (n = 175, 2 experiments). ***D,*** Bar histogram summarizing the amplitude ± SEM of RAP responses in the experiment shown in C. Statistical significance was calculated by a one-way ANOVA for repeated measures followed by Bonferroni post-hoc test. ***E,*** Average ± SEM Fura2 ratio of cold-evoked responses in CR#1 HEK293 cells in control and 30 μM RAP (n = 98). ***F,*** Bar histogram summarizing the effect of vehicle or 1, 10, 30 μM rapamycin on cold-evoked responses. Individual cold responses have been normalized to the amplitude during the first cold ramp (71 to 92 cells per condition). Ratios > 20 have been excluded from the analysis. Statistical comparison with respect to vehicle; one-way ANOVA followed by Bonferroni post-hoc test.

RAP has been shown to activate intracellular Ca^2+^-permeable channels, including TRPML1 at the membrane of lysosomes (Zhang et al., 2019), and ryanodine receptors in the endoplasmic reticulum (Brillantes et al., 1994; Lombardi et al., 2017). In order to corroborate that calcium responses evoked by RAP were due to calcium influx from the outside (presumably through TRPM8 channels) and not by calcium release from intracellular stores, the effect of RAP was explored in the control (2.4 mM Ca^2+^) and the calcium-free extracellular solution. During calcium imaging experiments, RAP (30 µM) evoked clear responses in the presence of extracellular calcium, whereas, when calcium was removed, RAP-evoked responses were absent (Fig. 1C-D).

### Rapamycin sensitizes TRPM8 channels to cold temperatures

TRPM8 agonists potentiate responses to cold (Voets et al., 2004; Mälkiä et al., 2007). This was tested in CR#1 cells expressing rat TRPM8, using a double pulse protocol. As shown in Figures 1E-F, cold-evoked responses were strongly potentiated in the presence of 1 to 30 μM RAP, compared to the response obtained in vehicle.

### Rapamycin activates different TRPM8 orthologs including human

Next, we studied the sensitivity of other TRPM8 orthologues to RAP. Transfected HEK293 cells expressing mouse TRPM8 were activated by a cooling ramp, by subsequent application of RAP (30 µM) and by menthol (30 µM) (Fig. 2A, blue trace). On average, RAP- and menthol-evoked responses were similar in amplitude and significantly smaller when compared to cold-evoked responses (Cold= 0.96 ± 0.08, RAP= 0.40 ± 0.03, menthol= 0.47 ± 0.03, n= 97) (Fig. 2B). In the presence of RAP or menthol, cold-evoked responses were strongly potentiated (Cold= 0.96 ± 0.08 *vs* RAP + Cold= 2.96 ± 0.13 *vs* Menthol + Cold= 2.43 ± 0.12) (Fig. 2B). Additionally, there was a subset of mTRPM8 transfected (i.e. GFP+) cells that did not respond to cold in control conditions but responded to cold in the presence of RAP or menthol (Fig. 2A, green trace). Cells not expressing mTRPM8 (i.e. those not responding to cold in the presence menthol) did not show changes in intracellular Ca^2+^ levels in the presence of RAP either (Fig. 2A, black trace).

**Figure 2.**
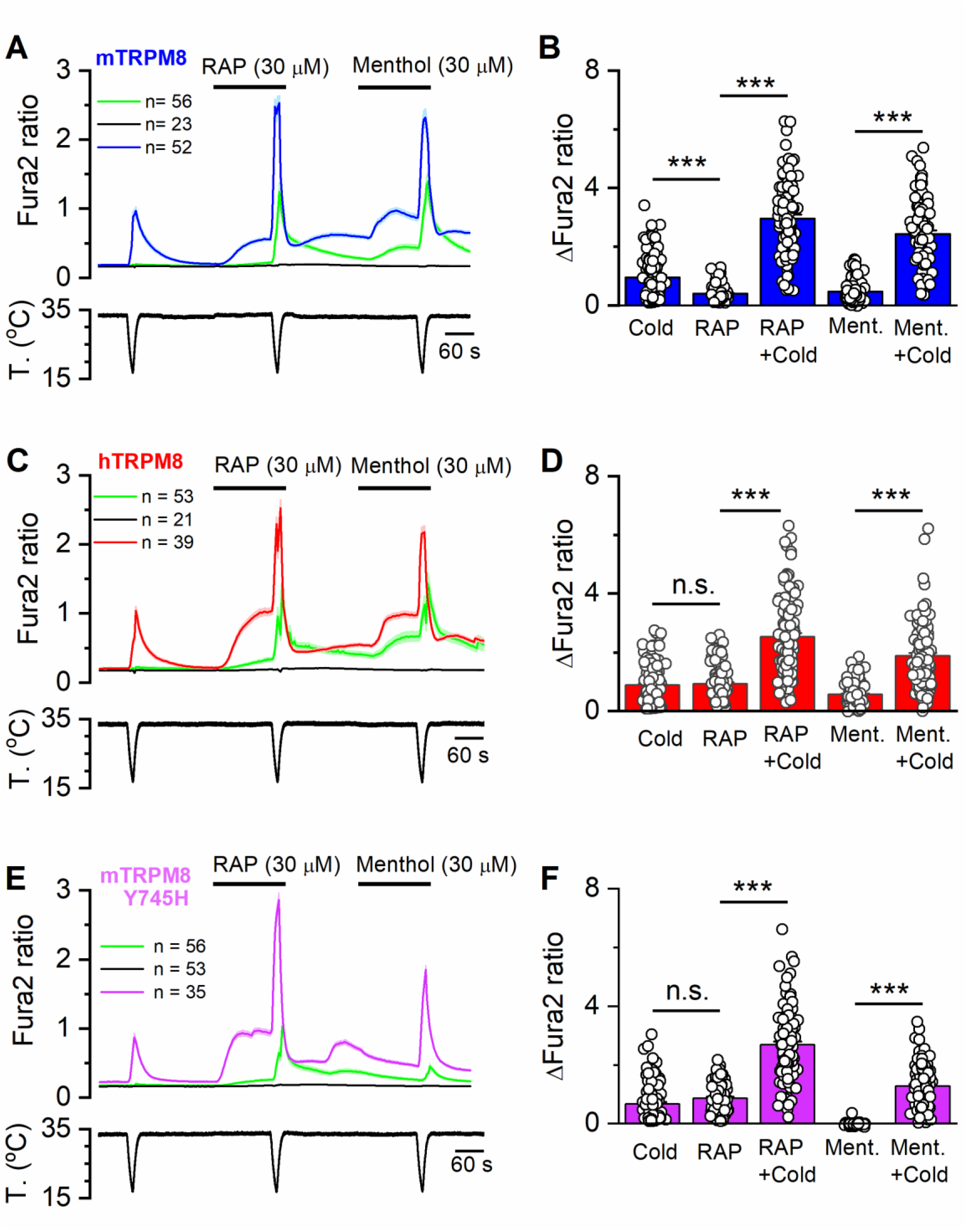
Rapamycin activates mouse and human TRPM8 orthologs and the menthol-insensitive mutant. ***A***, Average ± SEM Fura2 ratio time course in HEK293 cells expressing mouse TRPM8. Three different behaviors were observed; cells which responded to a first cold ramp in control conditions (blue trace), cells which did not respond to this cold ramp in control conditions but responded to cold in the presence of RAP or menthol (green trace), and cells which did not show any response to the applied stimuli (black trace). ***B***, Bar histogram summarizing the individual and mean responses of the cold-sensitive HEK293 cells (n = 97, 3 experiments. Statistical significance was calculated by a one-way ANOVA followed by Bonferroni post-test. ***C,*** Average ± SEM Fura2 ratio time course in HEK293 cells expressing human TRPM8. The different traces represented the three behaviours described in A. ***D,*** Bar histogram summarizing the responses of the cold-sensitive HEK293 cells (n = 115) transfected with human TRPM8. Statistical significance was calculated by a one-way ANOVA followed by Bonferroni post-test. ***E,*** Average ± SEM Fura2 ratio time course in HEK293 cells expressing mouse TRPM8-Y745H. The different traces represented the three behaviours described in A. ***F,*** Bar histogram summarizing the responses of the cold-sensitive HEK293 cells (n = 106, 3 experiments) transfected with human TRPM8. Statistical significance was calculated by a one-way ANOVA followed by Bonferroni post-test.

To verify that RAP-evoked calcium responses were mediated by TRPM8, we tested the effect of AMTB (10 μM), a selective TRPM8 antagonist (Lashinger et al., 2008). As shown in Figure S1, in the presence of AMTB, RAP-evoked calcium responses were strongly suppressed (mean amplitude 0.84 ± 0.07 *versus* 0.11 ± 0.11, p < 0.001, n = 69). As expected, cold-evoked calcium responses were also blocked by AMTB.

Next, we explored the sensitivity of human TRPM8 to RAP. In HEK293 cells transiently expressing hTRPM8, RAP (30 µM) evoked a robust response which was similar in amplitude to the cold-evoked response and significantly larger than the menthol-induced response at the same concentration (cold = 0.88 ± 0.06 *vs* Rapamycin= 0.93 ± 0.05 *vs* Menthol= 0.57 ± 0.04, n= 115) (Fig. 2C-D, red trace). hTRPM8-mediated cold-evoked responses were also strongly potentiated by RAP and, remarkably, this effect was stronger than the menthol-evoked potentiation (Cold= 0.88 ± 0.06 *vs* Rapamycin + Cold= 2.52 ± 0.12 *vs* Menthol + Cold= 1.88 ± 0.10, n= 115) (Fig. 2C, red trace). As in the case of mTRPM8, a subset of cold-insensitive cells in control conditions responded to cold in the presence of RAP and menthol (Fig. 2C green trace). Cells not expressing hTRPM8 did not show changes in intracellular Ca^2+^ levels in the presence of RAP (Fig. 2C, black trace).

In summary, RAP acts as an agonist of rat, mouse and human TRPM8 channels. Like other chemical agonists of TRPM8, RAP strongly potentiates the response to cold. In human TRPM8, this effect was more potent for rapamycin than for menthol, the canonical TRPM8 agonist.

### Rapamycin activates the menthol-insensitive TRPM8 mutant

Previously we reported that tacrolimus can activate the single-mutant TRPM8 (Y745H) that is insensitive to menthol (Bandell et al., 2006). The same result was obtained with RAP. As shown in figures 2E-F, RAP activated TRPM8-Y745H and potentiated cold-evoked responses.

### Everolimus does not activate but sensitizes responses to cold

Everolimus is a rapamycin analog, with similar immunosuppressant and antiproliferative properties by inhibition of the mTOR kinase (Houghton, 2010). Structurally, the only difference between both molecules is a 2-hydroxyethyl ether substituting for a hydroxy group on the cyclohexyl moiety. In HEK293 cells expressing mouse TRPM8, everolimus (30 µM) did not produce an increase in calcium but the responses to a cold stimulus were potentiated (Figure S2A-B). Similar results were obtained in cells transfected with human TRPM8 (Figure S2C-D). This behavior contrasts with the effects of RAP, tacrolimus (Arcas et al., 2019) or menthol, suggesting that everolimus is a weaker TRPM8 agonist.

### Effects of rapamycin on other thermoTRP channels

We and others have shown weak effects of tacrolimus on recombinant TRPA1 channels (Arcas et al., 2019; Kita et al., 2019). Thus, we tested the effects of RAP on human TRPA1. As shown in Figure S3A, RAP (30 µM) activated human TRPA1, with a slowly rising [Ca^2+^]_i_ response. The activating effects of RAP on TRPA1 were modest, compared to those produced by its canonical agonist AITC (Figs. S3A-B). Only around 60 % of the cells that responded to AITC were activated by RAP. In contrast, no activating effect of RAP was observed on cells expressing human TRPV1 (Fig. 3A) or rat TRPV1 (not shown), which responded readily to capsaicin.

**Figure 3.**
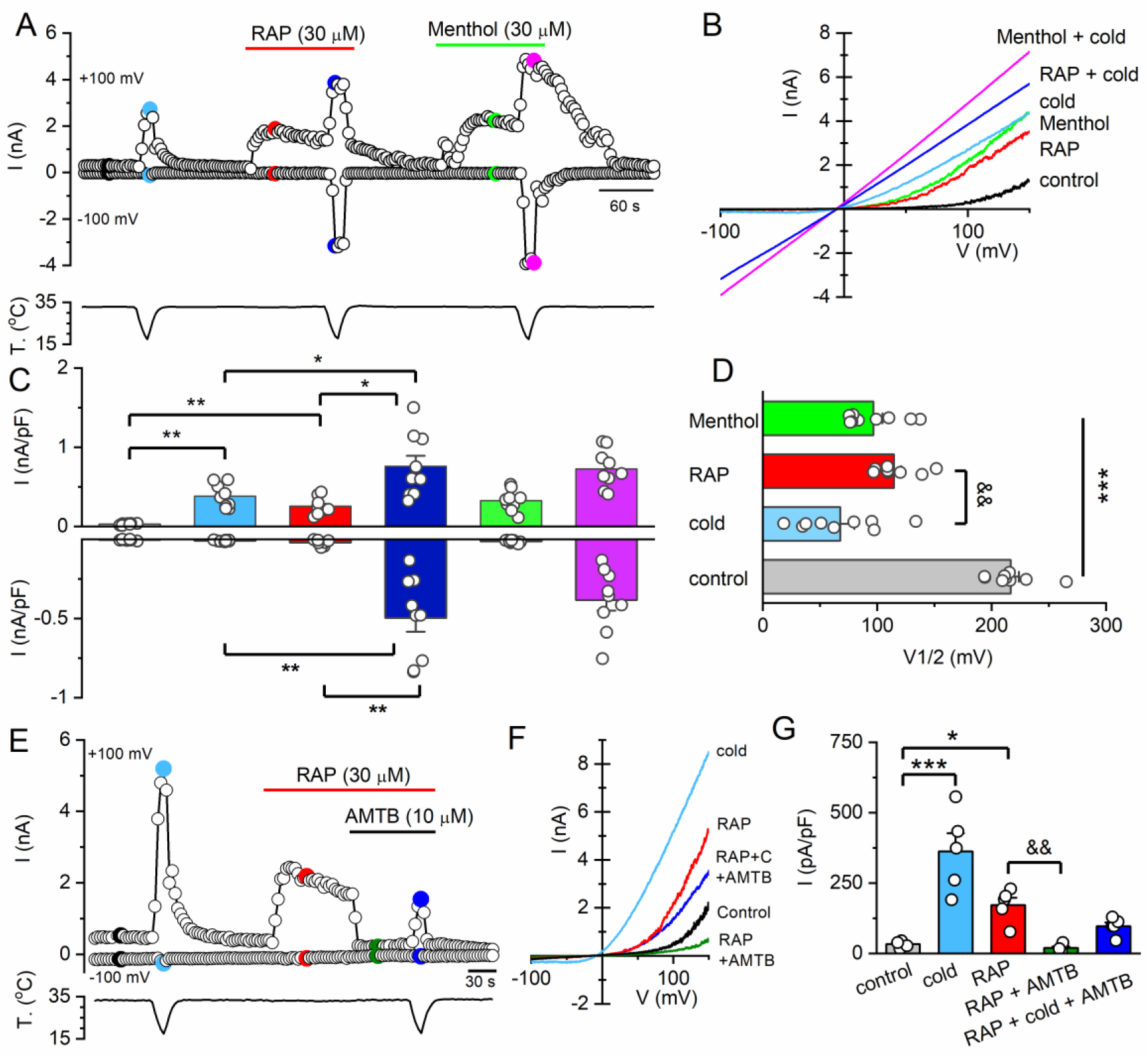
Rapamycin activates TRPM8 currents. ***A,*** Representative time course of whole-cell current at −100 and +100 mV in a HEK293 cell transiently transfected with mTRPM8 during the sequential application of different agonists. The bottom trace shows the simultaneous recording of the bath temperature during the experiment. ***B,*** Current-voltage (I-V) relationship obtained by a 400 ms voltage ramp from −100 to +150 mV during the experiment shown in A. The color of individual traces matches the color at each particular time point in A. ***C,*** Bar histogram of individual and mean ± SEM current density values at −100 and +100 mV evoked by the different stimuli shown in A, with the same color code (n = 9 cells, 2 experiments). Statistical differences were evaluated by a one-way ANOVA, followed by Bonferroni’s post-hoc test. ***D,*** Individual and mean ± SEM V_1/2_ values calculated from fitting individual I-V curves to a linearized Boltzmann equation. Statistical differences were evaluated by a one-way ANOVA, followed by a Bonferronís post-hoc test. ***E,*** Representative whole-cell current at −100 and +100 mV in a HEK293 cell transiently transfected with mTRPM8 during a protocol in which the effect of AMTB on rapamycin response was explored. Bottom trace corresponds to the simultaneous recording of bath temperature during the experiment. ***F,*** Current-voltage relationship (I-V) obtained with a 400 ms voltage ramp from −100 to +150 mV of the responses plotted in E. The color of the I-V curves matches the colored time points in E. ***G***, Bar histogram of individual and mean ± SEM current density values at +100 mV evoked by the different stimuli shown in E (n = 5 cells, 1 experiment). Statistical differences were evaluated by a one-way ANOVA, followed by Bonferroni’s post-hoc test. Asterisks are used for comparing the effect of the different stimuli with respect to control (*, p-value< 0.05; **, p-value< 0.01; ***, p-value< 0.001) whereas the & symbol is used for the comparison between two different stimuli.

### Rapamycin activates TRPM8 currents

Next, we studied the effect of RAP on TRPM8 channels by electrophysiological techniques. In whole-cell patch-clamp recordings, application of 30 µM RAP consistently activated whole-cell currents in HEK293 cells expressing mouse TRPM8 (Fig. 3A). The current-voltage relationship of the RAP-activated current showed strong outward rectification and a reversal potential close to 0 mV, in line with the known properties of TRPM8 channels (Voets et al., 2004) (Fig. 3B). These results confirmed the findings obtained in calcium imaging experiments.

Cold-evoked inward currents, measured at −100 mV, were strongly potentiated in the presence of 30 µM RAP (Cold= −10.2 ± 1.3 pA/pF *vs* RAP + Cold= - 497.3 ± 87.1 pA/pF, p<0.001, n = 9), a similar effect than that produced by menthol at the same concentration (Menthol = −13.8 pA/pF *vs* Menthol + Cold= −384.3 ± 66 pA/pF, p<0.001, n= 9). A similar potentiation of cold-evoked currents was observed when measured at +100 mV: (Cold = 381.5 ± 49.5 pA/pF *vs* RAP + Cold= 758.5 ± 134.5 pA/pF) and (Menthol = 326.8 ± 44.1 *vs* Menthol + Cold= 726.7 ± 80.2 pA/pF, n= 9). Moreover, RAP and menthol at the same concentration (30 µM) evoked outward currents of similar amplitude (RAP = 253.6 ± 34.9 pA/pF *vs* Menthol = 326.8 ± 44.1 pA/pF, p=0.32, n= 9) suggesting that the potency of both compounds as a TRPM8 agonist is similar. A summary of these results is shown in Figure 3C.

To analyze the effects of agonists on voltage-dependent activation of TRPM8, we estimated the V_1/2_ values in the presence of cold, RAP and menthol by fitting the traces derived from the voltage ramps with a Boltzmann-linear function (see Material and Methods) and compared them to the V_1/2_ value obtained in control conditions at 33 °C (Fig. 3D). The three agonists produced a notable shift of V_1/2_ toward more negative membrane potentials: RAP (ΔV_1/2_ = −102.1 ± 5.2 mV) and Menthol (ΔV_1/2_= −120.3 ± 7.6 mV) had a similar effect, while cold led to a more pronounced shift (ΔV_1/2_= −148.8 ± 6.6 mV), suggesting that it is a stronger agonist. The application of lower concentrations of RAP (3 and 10 μM) induced smaller currents and smaller shifts in V_1/2_ (Figure S4). These results, corroborate that RAP, as other TRPM8 agonists, activates TRPM8 by a shift in the activation curve towards physiologically relevant membrane potentials.

To confirm the agonism of RAP on TRPM8, we tested the effect of AMTB, a selective antagonist of TRPM8 channels (Lashinger et al., 2008). As shown in Figure 4E-F, AMTB fully suppressed TRPM8 currents evoked by RAP (RAP= 171.6 ± 26.2 pA/pF *vs* RAP + AMTB = 19.8 ± 4.9 pA/pF) (n= 5, p< 0.01). AMTB, at the concentration used (10 µM), also blocked the voltage-dependent activation of TRPM8 at the baseline temperature of 33 °C (not shown), although it did not totally block the response to cold in the presence of RAP (Fig. 3E-G).

**Figure 4.**
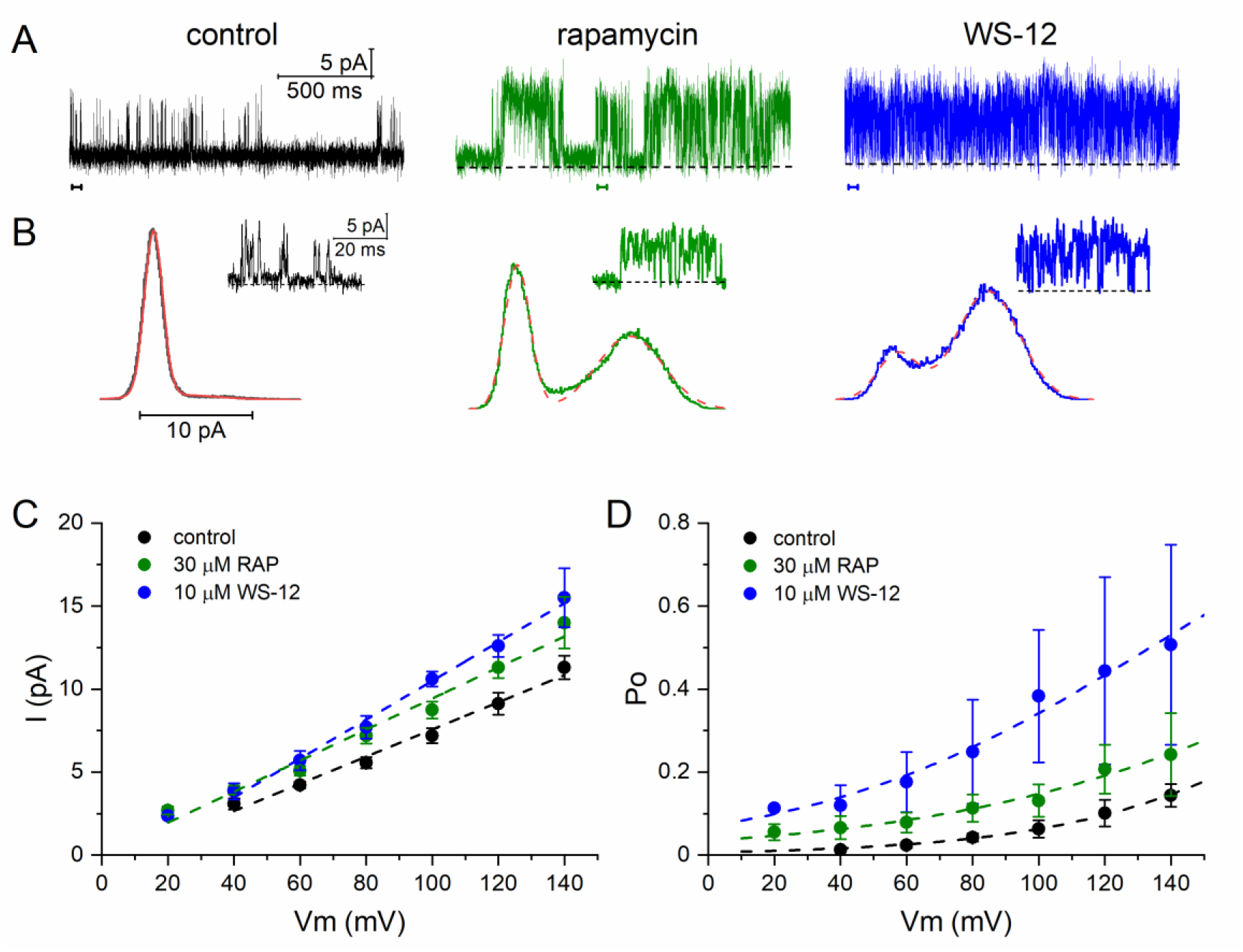
Rapamycin activates single TRPM8 channels. ***A,*** Single channel recordings at +100 mV in the cell-attached configuration in HEK293 cells transfected with mTRPM8. The same patch in control (i.e. high K^+^), 30 μM rapamycin and 10 μM WS-12. The colored ticks mark the expanded traces shown below. ***B,*** All-point histograms of current amplitudes for the different recording conditions. The histograms have been fitted with the sum of two Gaussians (red line). ***C,*** Single channel amplitudes of individual patches at different membrane potentials. The dotted lines represent linear fits to the data. ***D,*** Open probability of channel current at different potentials. The dotted lines are fits to the Boltzmann function.

### Gating of TRPM8 channels by rapamycin

To characterize the effect of RAP on TRPM8 channels at the single channel level, we recorded currents in the cell-attached configuration at room temperature (∼25 °C) in calcium-free, high potassium, extracellular solution (see Materials and Methods). At a fixed voltage (e.g. +100 mV), bath application of 30 μM RAP increased channel openings, detected as an increase in the peak current amplitude in the all-point histogram (Fig. 4A). After washing, the application of 10 µM WS-12 (produced openings of similar amplitude. The single channel conductance, estimated from the slope of the I-V relationship of well-resolved openings obtained at different positive membrane potentials was 98.8 ± 5.9 pS (n = 14) in RAP (Fig. 4C), very similar to the conductance estimate obtained during WS-12 application; 98.6 ± 10.6 pS (n = 5). Reversals potentials measured at the abscissa intercept of current amplitude were 3.2 ± 3.2 mV in RAP and −3.4 ± 5.3 mV in WS-12, values consistent with the non-selective nature of channel (Almaraz et al., 2014; McKemy et al., 2002).

The probability of channel opening induced by RAP was voltage-dependent, increasing at depolarized potentials (Fig. 4C), similar to the effect of WS-12, or other agonists (Zakharian et al., 2010). In the few cases where the patch contained a single channel, the increase in Po was explained by an increase in mean open time and a decrease in mean close time (not shown). Overall, these results indicate that RAP activates channels with biophysical properties similar to those induced by WS-12, a canonical TRPM8 agonist.

### Rapamycin activates TRPM8 in cold-sensitive DRG neurons

To study the effects of RAP on TRPM8-expressing sensory neurons, we performed calcium imaging experiments in DRG cultures from TRPM8^BAC-EYFP^ mice (Morenilla-Palao et al., 2014). TRPM8-expressing neurons were identified by their EYFP expression (Fig. 5A-B). Nearly all EYFP(+) neurons, 97% (33 out of 34), responded to a cooling ramp and 94.1% (32 out of 34) also responded to RAP (30 µM) application (Fig. 5C). In contrast, only a small percentage of EYFP(−) neurons were activated by 30 µM RAP (2 out of 236), although they showed normal responses to 30 mM KCl. In agreement with results obtained with recombinant channels, the amplitude of RAP-evoked responses was somewhat smaller than cold-evoked responses (RAP = 0.69 ± 0.06 *vs* cold = 0.89 ± 0.07) (n= 32, p< 0.05). Notably, their respective amplitudes correlated strongly in individual neurons (r^2^= 0.76) (Fig. 5E). The co-application of a cooling stimulus in the presence of RAP produced a marked potentiation in the amplitude of the cold-evoked response (Cold= 0.89 ± 0.07 *vs* RAP + Cold= 1.31 ± 0.09) (n= 32, p<0.001) (Fig. 5D).

**Figure 5.**
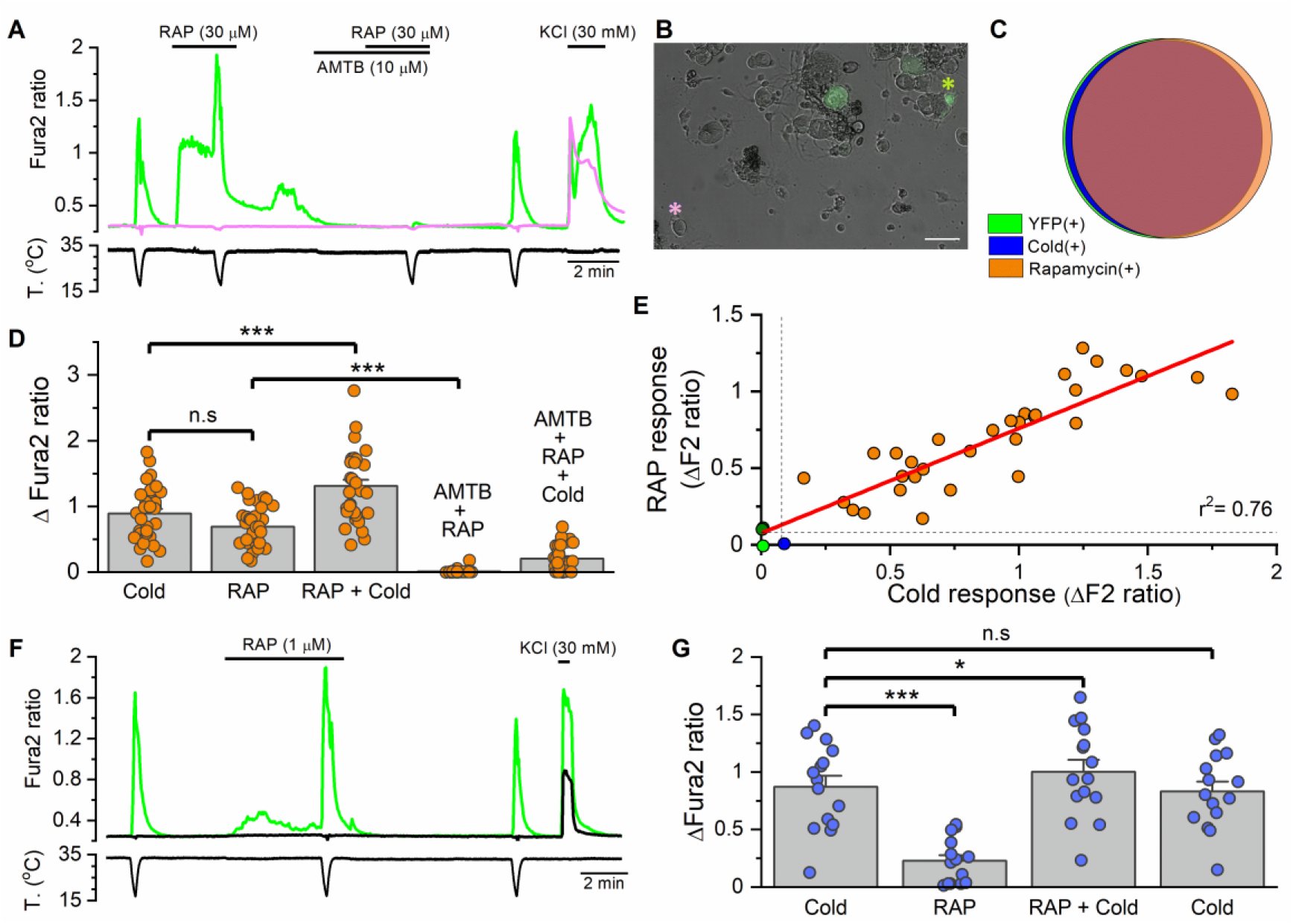
Rapamycin activates cold-sensitive neurons and potentiates their cold-evoked response. ***A*** Representative Fura2 fluorescence measurement from cultured DRG neurons of TRPM8^BAC-EYFP^ mice. A cold-sensitive and EYFP(+) neuron (green trace) and a EYFP(−) neuron (pink trace) are displayed. Bottom trace corresponds to the simultaneous recording of bath temperature. ***B,*** Superimposed transmitted and EYFP fluorescence images from DRG cultured neurons of a TRPM8^BAC-EYFP^ mouse. Asterisks correspond to the neurons whose traces are shown in A. The other two fluorescent neurons in the field had a baseline calcium level higher than 0.8 and were not included in the analysis. The calibration bar is 100 µm. ***C,*** Venn diagram showing the very strong overlap between EYFP(+) neurons (n= 34), cold-sensitive neurons (n= 33) and rapamycin-sensitive neurons (n= 32). Only 2 out of 236 EYFP(−) neurons were rapamycin-sensitive. ***D,*** Bar histogram of individual and averaged amplitude responses observed in EYFP(+) rapamycin-sensitive neurons (n = 32 neurons, 6 experiments). Statistical significance was assessed by a one-way ANOVA for repeated measures followed by Bonferroni’s post-hoc test. ***E,*** Correlation between amplitude of cold- and rapamycin-evoked responses in individual EYFP(+) DRG neurons (n= 34). Orange circles represent neurons that responded to rapamycin and cold, dark blue point represents the neuron which responded to cold but not to rapamycin (note the very small response to cold), the green point represents the single EYFP(+) neuron which did not respond to either stimulus and the olive points represent the only two EYFP(−) neurons which respond to rapamycin (overlapped). Dotted lines delimited the threshold that was considered as a response (ΔF2 ratio= 0.08). ***F,*** Time course of Fura2 ratio of a cold-sensitive EYFP(+) neuron (green trace) in which rapamycin (1 µM) evoked a substantially increase in intracellular calcium levels, and a cold-insensitive EYFP(−) neuron (black trace). Bottom trace corresponds to the simultaneous recording of bath temperature. ***G,*** Bar histogram summarizing the effect of rapamycin (1 µM) on cold-sensitive neurons (n = 15 neurons, 3 experiments). Statistical significance was assessed by a one-way ANOVA for repeated measures followed by Dunnett post-hoc test.

To explore the TRPM8-dependence of these responses, we studied a second RAP response in the presence of AMTB (10 µM), a specific blocker of TRPM8 channels (Lashinger et al., 2008). RAP responses were totally abolished in the presence of AMTB (RAP= 0.69 ± 0.06 *vs* RAP + AMTB= 0.01 ± 0.005) (n=32, p<0.001), confirming the absolute TRPM8 dependence of RAP responses in cold-sensitive DRG neurons (Fig. 5A, 5D). The cold-evoked response in the presence of RAP was also strongly reduced in the presence of this antagonist, and cold-sensitivity recovered after AMTB wash out (Fig. 5A, 5D).

Next, we examined the effect of a lower concentration of RAP (1 µM) on the thermal sensitivity of TRPM8-expressing DRG neurons (Fig. 5F). RAP 1 µM activated 67% (10 out of 15) of the cold-sensitive, and EYFP(+), neurons tested. Responses were of low amplitude (Fig. 5G). However, at this lower concentration, RAP significantly potentiates the cold-evoked response in a reversible manner (cold= 0.87 ± 0.09 *vs* RAP + cold= 1.0 ± 0.1) (n= 15, p<0.01) (Fig. 5F-G).

Collectively, these results indicated that RAP activates cold-sensitive TRPM8-expressing DRG neurons in a highly specific manner. RAP, even at low concentrations, strongly potentiates cold-evoked response of these neurons.

### Rapamycin responses in DRG neurons are mediated by TRPM8

To demonstrate the dependence of RAP responses on TRPM8, we took advantage of a transgenic mouse line (*Trpm8^EGFPf^*) expressing EGFPf from the *Trpm8* locus (Dhaka et al., 2008). Heterozygous mice (*TRPM8^EGFPf/+^*) in this line express one functional copy of TRPM8 and one copy of EGFPf, while the homozygous littermates (*TRPM8^EGFPf/EGFPf^*) are KO for TRPM8. In *TRPM8^EGFPf/+^* DRG cultures, all EGFPf(+) neurons (18 out of 18) responded to a cold ramp in control conditions and to menthol (100 µM), confirming the expression of TRPM8. They represented 16% of all the neurons in the culture. The majority of these TRPM8-expressing neurons, 83.3% (15 out of 18), also responded to RAP (30 µM) (Fig. 6A). In contrast, only 3 out of 112 EGFPf(−) neurons showed a response to RAP (Fig. 6B), a significantly lower fraction compared to the EGFPf(+) population (p<0.001, Z test). Cold- and RAP-induced responses were similar in amplitude (Cold= 0.98 ± 0.12 *vs* RAP = 0.73 ± 0.09) (n = 15, n.s.) and their amplitudes were correlated in individual neurons (r^2^= 0.57) (Fig. 6D). RAP was also effective in potentiating the cold-evoked response (Cold = 0.98 ± 0.12 *vs* RAP + Cold = 1.8 ± 0.2, n= 15, p<0.001) (Fig. 6C). This potentiating effect for RAP (30 µM) on cold-evoked responses was similar to that observed with menthol (100 µM) (Menthol + Cold 1.53 ± 0.16, n= 15, n.s.) (Fig. 6C).

**Figure 6.**
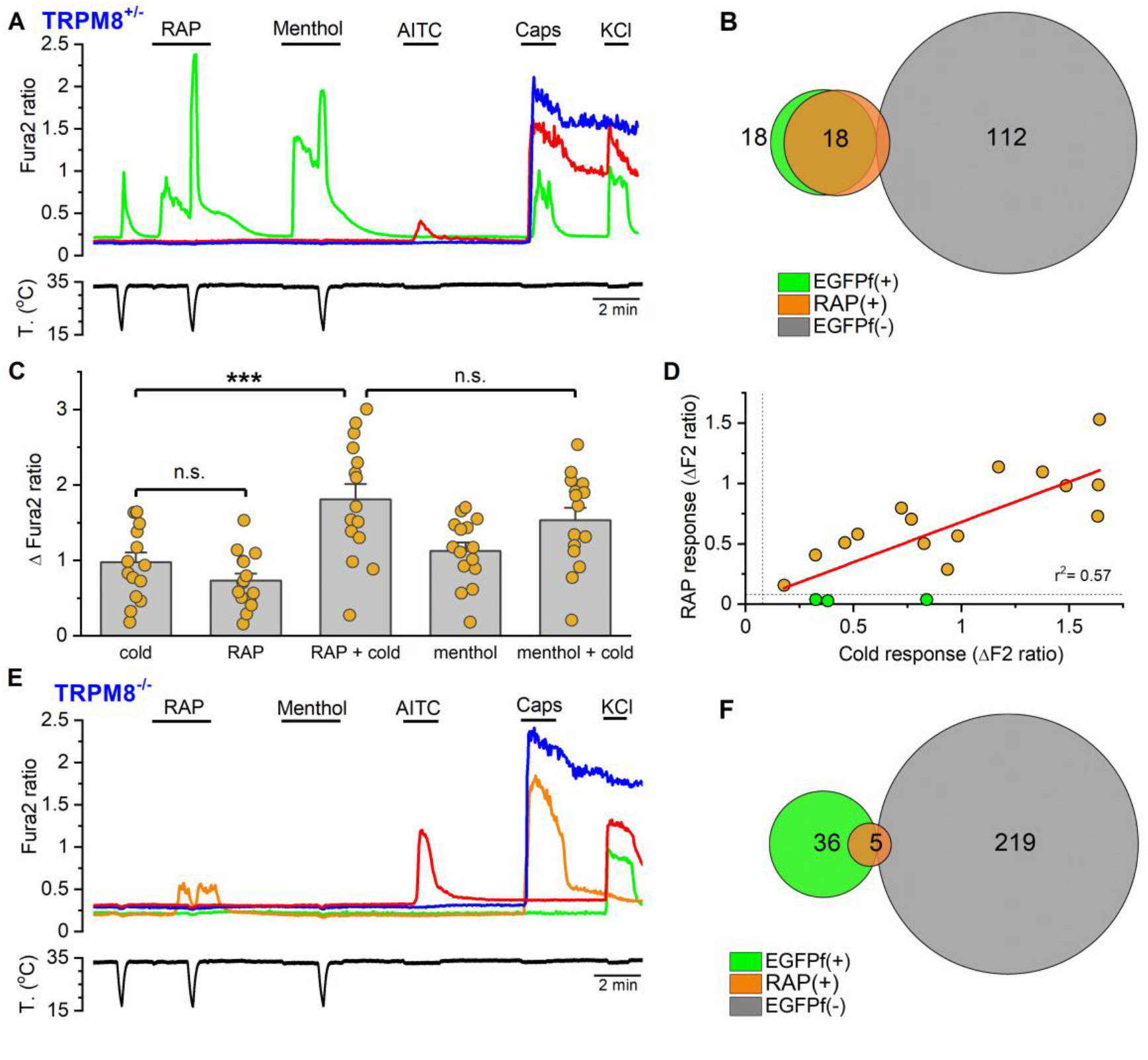
TRPM8 is the principal determinant of rapamycin responses in mouse DRG neurons. ***A,*** Representative traces of Fura2 ratio fluorescence in a TRPM8^EGFPf/+^ DRG culture. Consecutive applications of cold, rapamycin (30 µM), menthol (100 µM), AITC (100 µM), capsaicin (100 nM) and high K^+^ (30 mM) were used to phenotype each neuron. The green trace corresponds to a EGFPf(+) neuron which responds to cold, rapamycin, menthol and capsaicin. The other two traces (blue and red) correspond to EGFP(−) neurons. Bottom trace shows the simultaneous recording of bath temperature. ***B,*** Venn diagram summarizing the responses to rapamycin in EGFPf(+) and EGFPf(−) neurons in TRPM8^EGFPf/+^ DRG cultures. ***C***, Bar histograms of individual and mean ± SEM amplitude of the responses to different agonists in EGFPf(+) rapamycin-sensitive neurons (n = 15, 4 experiments). Statistical significance was analyzed by a one-way ANOVA for repeated measures followed by a Bonferroni’s post-hoc test. ***D*,** Correlation between the amplitude of the cold-evoked response and the rapamycin response (n= 15). The horizontal dotted line marks the threshold level for rapamycin sensitivity. Green circles represent the three EGFPf(+), cold-sensitive, rapamycin-insensitive neurons recorded. ***E*,** Representative traces of Fura2 ratios in cultured neurons from Trpm8 KO mouse. The protocol was the same as in A. Orange trace corresponds to a EGFPf(+) neuron that was rapamycin-sensitive. Green trace represents a EGFPf(+) neuron which was not sensitive to any of the agonists tested and the red and blue traces are examples of two EGFPf(−) neurons. Bottom trace corresponds to the simultaneous recording of the bath temperature during the experiment. ***F*,** Venn diagram summarizing the responses to rapamycin in EGFPf(+) and EGFPf(−) neurons in TRPM8^EGFPf/ EGFPf^ DRG cultures (8 experiments).

Confirming previous findings (Arcas et al., 2019; Dhaka et al., 2007), in DRGs from homozygous mice (*Trpm8^EGFPf/EGFPf^*) the responses to menthol and cold were abrogated; none of the 36 EGFPf(+) recorded neurons responded to menthol and only 2 of them responded to cold. Rapamycin responses were also very infrequent in these cultures. Only 5 out of 257 (1.9 %) neurons responded to RAP, 3 were EGFPf(+) and 2 were EGFPf(−) (Fig. 6E-F). The decrease in the percentage of RAP responses in EGFPf(+) neurons was highly significant (p<0.001, Z test). Interestingly, the RAP responses in these three EGFPf(+) neurons were maintained during the time of application and decreased during the cooling ramp (Fig. 6E), while the responses of EGFPf(−) neurons were transient and smaller in amplitude (not shown). Two of these EGFPf(+) RAP sensitive neurons responded to capsaicin while the other was capsaicin-insensitive, AITC-insensitive, but was activated by cold and cold in the presence of menthol (not shown).

Altogether, the results obtained in *Trpm8* KO mice indicate that TRPM8 channels mediate the main excitatory action of RAP in DRG neurons.

### Rapamycin activates inward currents and elicits AP firing in TRPM8 cold thermoreceptors

Next, we investigated the effects of RAP on cold-sensitive DRG neurons by patch-clamp recordings in cultured neurons from TRPM8^BAC-EYFP^ mice. In the whole-cell, voltage-clamp configuration, a cooling ramp in control conditions activated a small inward current (Fig. 7A). Application of 30 μM RAP at 33 °C also activated a sustained, small inward current and strongly potentiated the cold-evoked response (Cold= −12.6 ± 3.5 pA/pF *vs* RAP + Cold= −50.7 ± 7.5 pA/pF) (n = 12, p<0.001). A summary of these results is shown in Figure 7C. In the presence of RAP, the apparent activation of the cold-evoked current shifted to warmer temperatures (Fig. 7B). The potentiated currents showed variable degrees of desensitization, likely reflecting the effects of elevated intracellular calcium on TRPM8 gating (Reid et al., 2002; Sarria et al., 2011). In a fraction of neurons, after the RAP challenge, we tested the application of menthol (30 μM), which produced similar effects (Figure S5).

**Figure 7.**
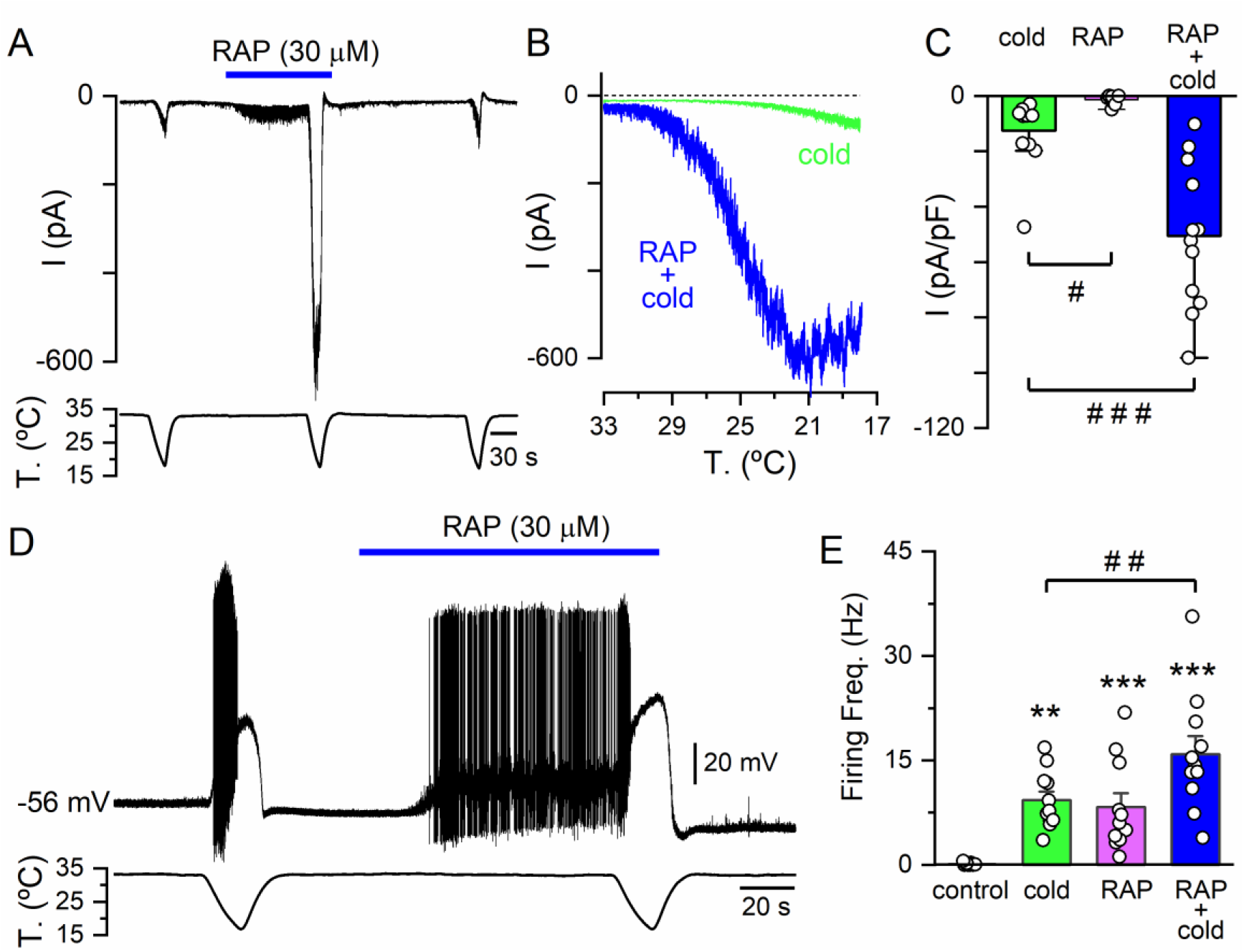
Rapamycin increases the excitability of cold-sensitive DRG neurons. ***A*,** Representative whole-cell recording in the voltage-clamp configuration (V_hold_= −60 mV) of a TRPM8-expressing, cold-sensitive DRG neuron identified by EYFP expression. Bottom trace corresponds to the simultaneous recording of bath temperature during the experiment. RAP activates a small inward current and potentiates the response to cold. ***B,*** Temperature dependence of the cold-evoked current in the same neuron in control conditions and in the presence of 30 μM RAP. ***C*,** Bar histogram of individual and mean peak inward current density values evoked by cold, RAP and cold in the presence of RAP (n = 12, 3 experiments). The statistical analysis consisted on a one-way ANOVA for repeated measures followed by Bonferroni’s post-hoc test. ***D,*** Representative recording of a cold-sensitive neuron in the whole-cell current-clamp configuration. Cold and RAP elicited the firing of action potentials. The combined application of cold and RAP led to faster firing, followed by a strong depolarization and the blockade of spikes. ***E,*** Bar histogram of individual and mean responses, measured as average firing frequency, during the different stimuli applied (n = 11, 3 experiments). Firing frequency for cold was the average from the first to the last spike during the cooling ramp. Firing frequency in control conditions was calculated during the 60 s previous to RAP application. RAP-evoked firing was calculated from the first spike during rapamycin application to the start of the cold ramp. The asterisks denote statistical differences with respect to control, and the # # differences between cold and cold + RAP, all calculated by a one-way ANOVA for repeated measures followed by Mann-Whitney post-hoc test.

In the current-clamp recording configuration, RAP induced the firing of action potentials in cold-sensitive neurons at a basal temperature of 33 °C in all the neurons tested (11 out of 11) (Fig. 7D-E). The cold- and RAP-evoked firing frequency was similar (Cold = 9.2 ± 1.2 Hz *vs* RAP = 8.3 ± 2.0 Hz, n = 11, p = 1). The combined application of cold and RAP strongly potentiated the firing produced by cold alone (Cold = 9.2 ± 1.2 Hz *vs* RAP + Cold= 15.9 ± 2.6 Hz, n= 11, p= 0.002) (Fig. 7E). The duration of firing during the cooling ramp in the presence of agonist is limited by the spike inactivation produced by the strong depolarization (Fig. 7D).

These results corroborate that RAP is able to trigger a depolarizing current in TRPM8(+) cold-sensitive DRG neurons, increasing their excitability to cold temperatures.

### Rapamycin activates cutaneous cold-sensitive fibers

To characterize the effects of RAP on cutaneous cold thermoreceptor endings, we used a mouse skin-nerve preparation of the hind leg (Arcas et al., 2019; Zimmermann et al., 2009), probing the corium surface of the skin with a miniature ice pellet, trying to find unimodal cold receptors (i.e. insensitive to mechanical stimuli) (Toro et al., 2015; Winter et al., 2017). We identified 5 fibers in the saphenous nerve with the aforesaid characteristics. These fibers were silent at the baseline temperature of 34-35 °C but were activated when cold solution was delivered to their isolated receptive field (Fig. 8A), with a mean cold threshold of 29.1 ± 0.9 °C (n = 5) (Fig. 8C). During application of RAP (30 μM) all five fibers became spontaneously active at the basal temperature of 34 °C. Their cold-evoked activity was clearly modified in the presence of RAP, shifting their cold threshold to warmer temperatures (mean temperature threshold displacement of 4.5 ± 0.9 °C, n = 5) (Fig. 8C) and shifted their stimulus response function to warmer temperatures (Fig. 8B), consistent with the results obtained for cold-sensitive DRG neurons. In agreement with the effects observed with tacrolimus (Arcas et al., 2019), the washout of RAP effects was only partial (Fig. 8C-D). Subsequent application of menthol (50 µM) produced a reactivation of firing in all the fibers tested. Collectively, these results indicate that RAP sensitizes a population of cutaneous TRPM8-expressing thermoreceptor endings to cold temperature

**Figure 8.**
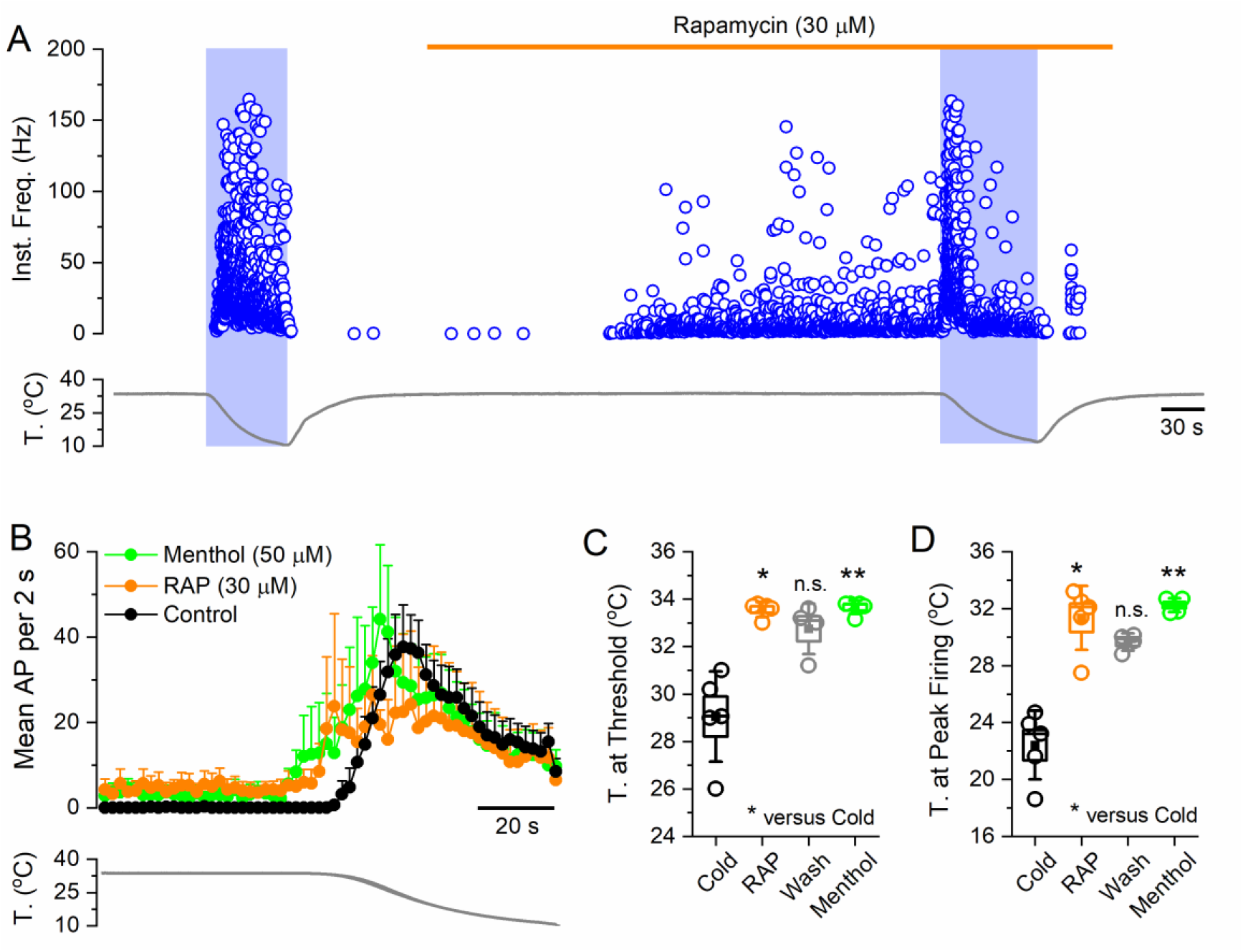
Rapamycin activates cutaneous cold-sensitive nerve endings. **A,** Representative recording showing the firing response of a single saphenous nerve cold fiber to a decrease in the temperature of the receptive field. The circles represent the instantaneous firing frequency of the fiber. **B,** Time course of the averaged cold-evoked response of cold fibers from C57BL/6J mice in control solution, in the presence of 30 μM RAP and in 50 μM menthol. Average discharge rates are represented in bins of 2 s (n = 5, 5 experiments). Bottom, The average temperature ramp for each of the datasets. **C,** Temperature threshold for activation of impulse discharge for the different experimental conditions. **D,** Temperature at the maximal discharge rate. Circles represent mean values (n=5). Boxes represent SEM. Error bars indicate standard deviation (SD). In C and D, statistical significance assessed by Kruskal-Wallis followed by Dunn’s test. The asterisks compare significance with respect to cold.

### Mathematical modelling of rapamycin effects on cold thermoreceptor activity

To obtain further insight into the effects of RAP on the excitability of cold-sensitive DRG neurons, we implemented the conductance-based mathematical model developed by Olivares et al. 2015 (Olivares et al., 2015). In addition to ion conductances needed for action potential generation, this model incorporates a depolarizing cold- and voltage-dependent current (i.e. TRPM8-like). The effects of RAP on TRPM8 were simulated as changes in the voltage-dependent activation (V_½_) (Voets et al., 2004). We explored the same set of thermosensitive DRG neurons as described in (Rivera et al., 2021), including low- and high-threshold neurons (Madrid et al., 2009). Resembling findings observed in culture, simulated neurons showed no spontaneous activity at rest but became vigorously active during cooling ramps (Fig. 9A). In current-clamp mode, in the presence of RAP the neuron starts firing at the basal temperature, increasing its peak firing frequency further during a second cooling ramp. As shown in Fig. 9B-C, changes in V_½_ produced graded changes in the temperature threshold and the average firing frequency of low- and high-threshold neurons.

**Figure 9.**
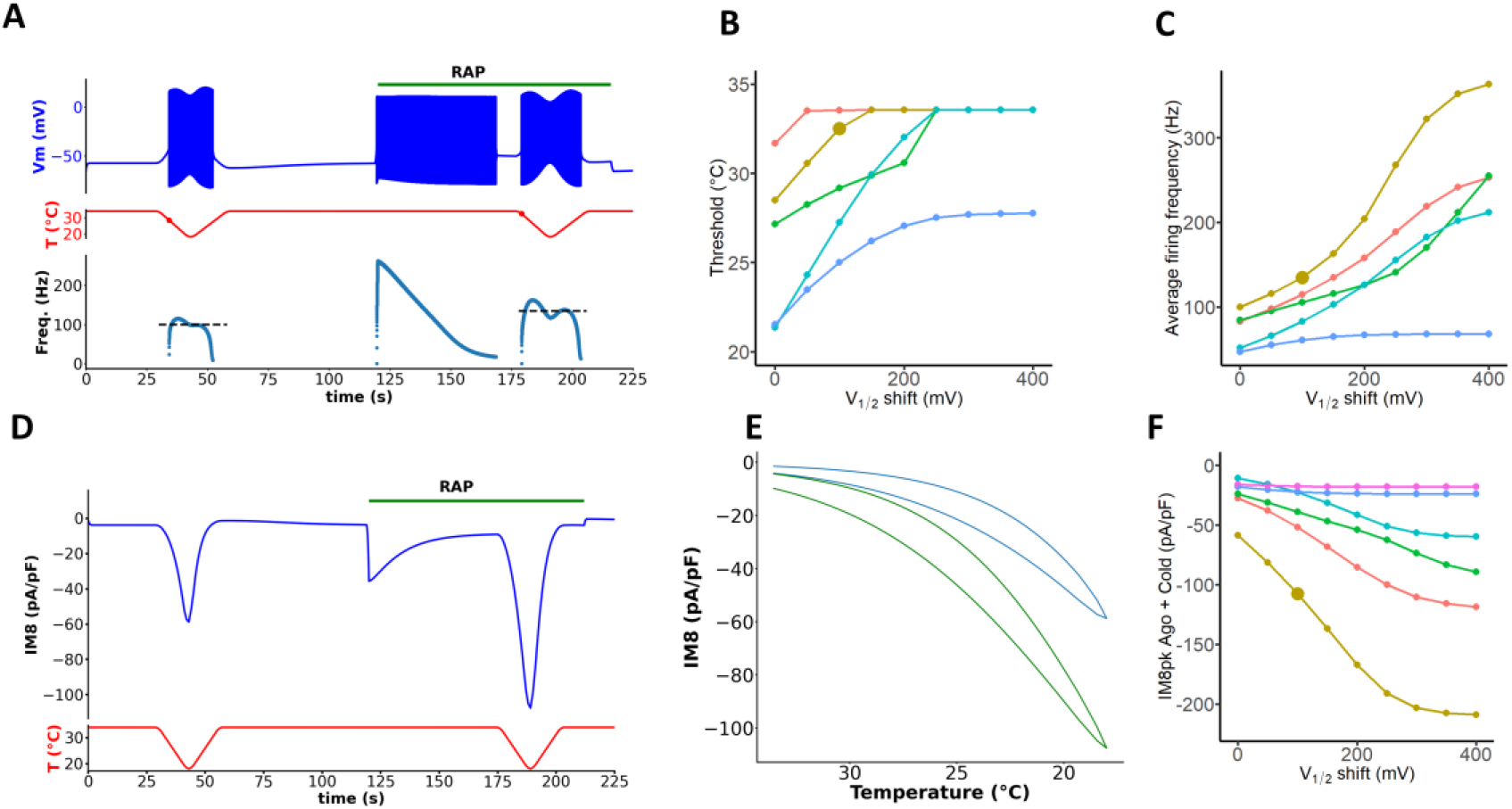
Mathematical modelling of rapamycin effects on cold thermoreceptor activity. **A.** Membrane potential (blue trace), temperature (red trace), and action potential firing frequency (blue dots), as a function of time. The green bar indicates the period of application of agonist, i.e. a change of −100 mV in V_1/2_ in this example. The red dots in the temperature trace indicate where the temperature threshold has been reached. The black dashed lines in the lower panel indicate the value of the average firing frequency, and the time range where it has been calculated (i.e. the duration of the temperature ramps). **B.** Firing temperature threshold as a function of the shift in V_1/2_. Each trace corresponds to a cell with different parameters, that have been selected in order to encompass a wide range of thresholds under control conditions. The larger dot corresponds to the cell and conditions simulated in A. **C.** Same as in B for the average firing frequency during cold stimulation. **D.** TRPM8 current density (blue trace) as a function of time, and changes in temperature (red trace) in voltage-clamp simulations. The green horizontal bar represents the application of agonist (i.e. a shift of −100 mV in V_1/2_ as in A). **E.** Current density as a function of temperature with (green trace) or without agonist (blue trace). **F.** Peak cold-evoked current as a function of the shift in V_1/2_ for the same cells as shown above with the corresponding colors. Values for D V_1/2_ = 0 correspond to the cold-evoked current in the absence of agonist. The magenta line corresponds to a neuron with very small TRPM8 currents that did not fire action potentials and thus is not shown in the current-clamp panels above.

In voltage-clamp mode, application of RAP at 34 °C produces a modest sustained inward current and a marked increase in peak inward current at the lowest temperature reached (i.e. 18 °C) (Fig. 9D). Similar to the experimental findings (not shown), the amplitude of the current as a function of temperature presents a strong hysteresis that is traversed with time in counterclockwise direction (Fig. 9E). The increase in cold activated-activated current as a function of RAP concentration, modelled as shifts in V½ (ΔV_1/2_ = 0 mV represents the condition before agonist application) in is shown in Figure 9F for six different DRG neurons.

In summary, the experimental effects of RAP on cold-thermoreceptor excitability are closely mimicked by effects of RAP on TRPM8 gating.

### Rapamycin application to the eye evokes tearing by a TRPM8-dependent mechanism

TRPM8 channels play a major role in basal tearing and blinking (Parra et al., 2010; Quallo et al., 2015), and TRPM8 agonists increase tearing and relieve the symptoms of dry eye (Arcas et al., 2019; Parra et al., 2010; Wirta et al., 2022; Yang et al., 2017).

We examined the effect of RAP solutions on tearing in anesthetized adult WT mice of both sexes. We applied a small drop of vehicle solution to one eye and measured the tearing after a rest period of 5 min. This was followed by application of 1% RAP to the other eye. The experimenter was blind to the solution applied to each eye, and the order of application was also randomized. As shown in Figure 10, in wildtype mice, RAP produced a significant increase in tearing compared with vehicle (p = 0.006; Mann-Whitney test). In contrast, in *Trpm8* KO mice application of RAP reduced tearing rate, and the response was similar to the application of vehicle (p = 0.27). These results indicate that TRPM8 channels mediate the effects of RAP on tearing.

**Figure 10.**
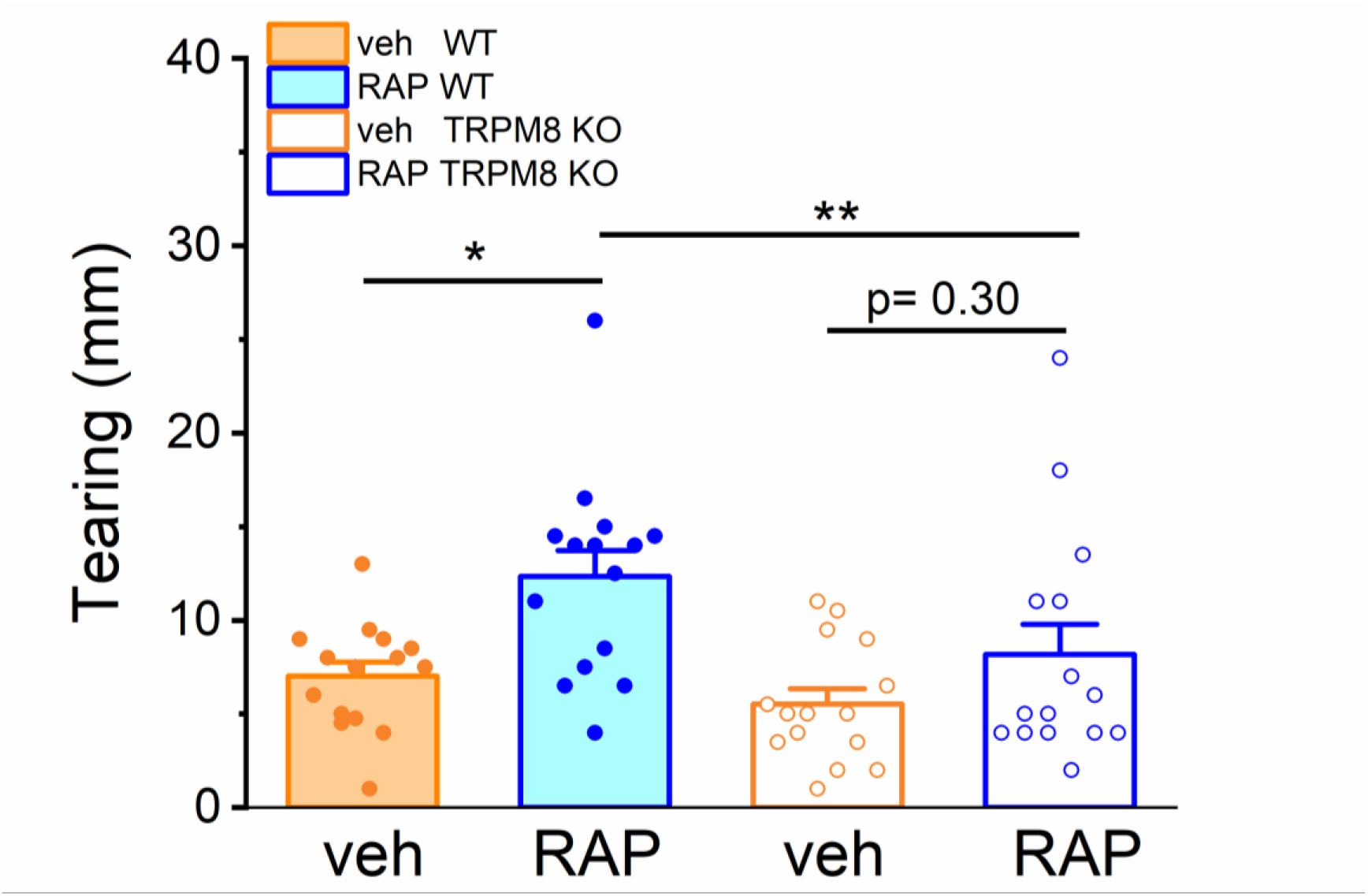
Rapamycin stimulates tearing by a TRPM8-dependent mechanism. Effect of topical solutions of RAP 1% or vehicle on tearing in mice. Tearing is represented in the box plots as the length of staining in the threads (in mm). Each dot corresponds to the tearing measured in an eye of an individual mouse (15 WT and 15 *Trpm8* KO). Histograms shows evoked tearing mean ± SEM (* p < 0.05, **p < 0.01. Kruskal-Wallis test).

## Discussion and Conclusions

Natural products represent a rich resource for modulators of different TRP channels (Julius, 2005; Meotti et al., 2014; Nilius & Appendino, 2011). In the case of TRPM8, menthol, a monoterpenoid extracted from mint leaves, was crucial for its discovery and functional characterization (McKemy et al., 2002; Peier et al., 2002; Voets et al., 2004). We found that RAP, a clinically relevant macrolide immunosuppressant produced by soil microorganisms, and its close analog everolimus, are novel agonists of TRPM8 channels. They share this property with another macrolide immunosuppressant, tacrolimus (Arcas et al., 2019), generalizing the structure of macrolide rings as chemical activators of TRPM8.

It is noteworthy that RAP and tacrolimus (Arcas et al., 2019) can both activate the menthol-insensitive Y745H mutant (Bandell et al., 2006). This result suggests that these macrolides could be binding to the channel at a different site. In this regard, a preliminary report by Toth et al., (https://www.researchgate.net/publication/354599833_Rapamycin_activates_TRPM8_via_novel_mechanisms) also reported activation of TRPM8 by RAP and other analog macrolides. In docking simulations, these authors identified possible RAP binding sites of TRPM8 independent of the menthol binding pocket. Additional work, including the screening of additional orthologs and mutagenesis, is currently underway to identify critical residues for RAP activation of TRPM8.

RAP potentiates cold-evoked responses by shifting the TRPM8 temperature threshold of activation in a dose-dependent manner, with a clear effect at low micromolar concentrations. Gating effects are similar to those reported for other Type I TRPM8 agonists, including TAC, characterized by the stabilization of the open channel state and a shift in the voltage dependence of activation towards more negative potentials (Janssens et al., 2016). RAP appears to be more potent than TAC, both in recombinant systems and in neurons. For example, in mouse DRG neurons, 1 µM RAP produced clear potentiating effects on cold-evoked responses, stronger than those observed with 10 µM TAC (Arcas et al., 2019). RAP-evoked responses are on par to those obtained with the same concentration of menthol, suggesting similar potency as a TRPM8 agonist. In summary, RAP activates a depolarizing inward current similar to the TRPM8-dependent I_Cold_ current, increasing the excitability of cold-sensitive DRG neurons.

### Specificity of RAP effects on TRPM8

A recent study found that RAP also activates Transient Receptor Potential channel mucolipin 1 (TRPML1) directly, at micromolar concentrations in human fibroblasts (Zhang et al., 2019). This is a lysosomal Ca^2+^-release channel, implicated in autophagy. However, multiple lines of evidence suggest that TRPML1 is not involved in the responses we observed. First, according to the authors, HEK293 cells (i.e. the heterologous expression system we used) are insensitive to RAP (Zhang et al., 2019). Second, our detailed pharmacological characterization of responses in wildtype, hemizygous and TRPM8 KO mice indicates that RAP activates TRPM8-expressing sensory neurons in a highly selective manner. Indeed, at the concentration tested, very few RAP-responsive cells did not have a TRPM8-like phenotype. In addition, TAC also activates TRPM8 (Arcas et al., 2019), but failed to activate TRPML1 (Zhang et al., 2019). Collectively, these results suggest that, in DRG neurons, the effects of RAP and other macrolides are mediated by TRPM8.

In contrast to the very selective effects of RAP on TRPM8-expressing mouse sensory neurons, RAP also produced a mild activation of human TRPA1 channels overexpressed in HEK293 cells. After topical application of RAP for the treatment of facial angiofibromas, patients reported significant skin irritation (Foster et al., 2012; Mutizwa et al., 2011), suggesting that TRPA1 activation may underlie this side effect.

### Rapamycin as an experimental tool

RAP has been used extensively in cell biology research and other investigations to induce heterodimerization of fusion proteins. This is due to RAP ability to bind with high affinity to FKBP12 and the FRB domain of mTOR, acting as a sort of cell-permeant molecular glue of protein fragments engineered to contain these two domains (Choi et al., 1996; Mangal et al., 2018). Our finding of RAP agonistic effect on TRPM8 was serendipitous. We were trying to reduce membrane PIP_2_ levels with pseudojanin, an enzymatic chimera of inositol polyphosphate 5-phosphatase type IV that depends on RAP dimerizing properties for activity (Hammond et al., 2012). Contrary to our expectations, addition of RAP to cells expressing pseudojanin did not inhibit TRPM8. Others have used this strategy successfully to investigate PIP_2_ depletion on TRPM8 (Varnai et al., 2006; Zhang, 2019). We suggest an explanation for these divergent results. In the study of Varnai, RAP was applied at low concentrations (100 nM) after maximal activation of TRPM8 by 500 μM menthol at room temperature. In contrast, we were working at 34 °C and used higher concentrations of RAP. It is possible that, under their experimental conditions, the agonist effect of RAP is masked by the depletion of PIP_2_ induced by the phosphatase. The results obtained in the study by Zhang are more difficult to compare with ours because he used preincubation of cells with 1 μM RAP, comparing TRPM8 current density with a separate control group.

### Unexplained effects of RAP

RAP administration has broad effects on the organism biology and some of them are still poorly understood. There is an intriguing connection between RAP administration and increased lifespan in multiple species, including mice (Bitto et al., 2016), but the mechanism is unknown. Remarkably, cold ambient temperature, the physiological agonist of TRPM8 and TRPA1 channels, also increases longevity in many poikilothermic species (Loeb & Northrop, 1916; Xiao et al., 2013). In homeotherms, like mice, the influence of ambient temperature on life span is more complex, due to physiological thermoregulatory mechanisms (Conti, 2008). Nevertheless, in mice with reduced core body temperature, life span is also extended (Conti et al., 2006). Intriguingly, TRPM8 deletion causes lipid accumulation and increased weight in mice (Reimundez et al., 2018), opposite to effects of RAP treatment (Bitto et al., 2016). Further studies are needed to establish a causal link between RAP treatment, TRPM8 activation and extended lifespan. Characterizing the effects of RAP treatment on lifespan in TRPM8 KO mice could shed light on the mechanisms involved in life extension.

A known side effect of RAP treatment is testicular toxicity, with impaired spermatogenesis and reduced testosterone levels (Rovira et al., 2012). TRPM8 is highly expressed in human and mouse sperm where it participates in different aspects of their physiology (Martinez-Lopez et al., 2011). In light of our findings, and the fact that testosterone is an activator of TRPM8 (Asuthkar et al., 2015), a possible connection between RAP and TRPM8 in the testis is worth exploring further.

### Therapeutic implications

TRPM8 channels play different roles in somatosensation, from sensing noxious cold (Knowlton et al., 2013), to participating in menthol (Liu et al., 2013) and cooling-mediated analgesia (Proudfoot et al., 2006), or relieving pruritus (Palkar et al., 2018). TRPM8 is also overexpressed in various types of cancers, including melanoma (Tsavaler et al., 2001). Due to their potential therapeutic benefit, identification of novel TRPM8 modulators (agonists and antagonists) has been in the agenda of many pharmaceutical companies and academic research groups (Izquierdo et al., 2021; Moran & Szallasi, 2017). Because of severe side effects, it is doubtful that systemic RAP will find an application as an anti-inflammatory agent. In contrast, topical applications could be beneficial for different conditions, including ocular pathologies and skin diseases. TRPM8 channels play an important role in the pathophysiology of dry eye disease (DED) (Parra et al., 2010), and several clinical trials support that TRPM8 agonists can relieve symptoms of ocular discomfort (Wirta et al., 2022; Yang et al., 2017). Our findings suggest that topical application of RAP could be a novel treatment for DED by activation of TRPM8. In fact, topical formulations of RAP are currently used for the treatment of various ocular disorders in humans and other animals. In particular, it is effective in the treatment of keratoconjunctivitis sicca, a very common disease in dogs, by stimulating tear production (Spatola et al., 2018). In this study, tear production was also increased in control animals, suggesting that the drug has direct lacrimostimulant properties, independently of its anti-inflammatory and immunosuppressive activity. In relation to skin pathologies, TRPM8 is expressed in keratinocytes (Bidaux et al., 2015; Denda et al., 2010) and human melanoma cells (Yamamura et al., 2008). Topical application of TRPM8 agonists accelerates epidermal permeability barrier recovery after injury (Denda et al., 2010). In cultured human melanoma G-361 cells, application of menthol produced a dose-dependent decrease in their viability (Yamamura et al., 2008). Inhibition of mTOR by rapamycin and its analogs continues to be explored as a viable therapy in various cancers (reviewed by Zou et al., 2020). Further preclinical studies are required to explore the link between TRPM8 modulation by RAP and tumor cell growth.

Our findings indicate that RAP, a drug already approved for clinical use, may be repurposed for the treatment of pathologies related to TRPM8 dysfunction, a concept that is called “drug repositioning” (Doan et al., 2011). This is an interesting strategy in drug discovery, with some important advantages compared to the *de novo* identification and development of active compounds. Since safety and pharmacokinetic profiles of repositioned candidates are already well stablished, it allows a dramatic reduction in development time and expense.

In summary, these findings identify RAP as a novel TRPM8 agonist and generalize the agonist effects of different macrolide immunosuppressant’s towards this polymodal ion channel.

## Author contributions

A significant fraction of this data is part of the PhD Thesis of the first author, José Miguel Arcas, defended at the Universidad Miguel Hernández in 2019, available on-line at https://dialnet.unirioja.es/servlet/tesis?codigo=221625. JMA, KO and JC performed patch-clamp recordings and their analysis, JMA, KO and SP performed calcium imaging recordings and their analysis, JFT and FP performed behavioral experiments, A Gonzalez performed skin nerve recordings and analysis, SS performed computer simulations, SS and EP performed single-channel recordings and their analysis, FT designed TRPM8 expression plasmids, AG and FV obtained funding and supervised the project, FV wrote the initial draft of the manuscript. All authors contributed to the writing and revision of the final version of the manuscript.

## VI. Acknowledgements

We thank Ardem Patapoutian and Ajay Dhaka for providing the Trpm8EGFPf mouse line, Patricio Orio for help with computer simulations and Juana Gallar for help with tearing measurements. M. Tora, E. Quintero, R. Torres and the SHARE service IN are acknowledged for excellent technical assistance. During the course of this work, JMA was supported by a predoctoral fellowships from the Spanish Ministry of Education and Science (SO). KO was supported by a fellowship from Generalitat Valenciana (GRISOLIA/2019/089). JC is recipient of a JAE Intro fellowship from CSIC (JAEINT_22_01043). SP was supported by Erasmus+ program of the European Union. The work was funded by Generalitat Valenciana (PROMETEO/2021/03), the International Center for Aging Research (ICAR; project ionRAPA) and the Spanish MCIU/AEI through projects PID2019-108194RB-I00/AEI/10.13039/501100011033, PID2022140961OB-100/AEI/10.13039/501100011033 and Centro de Excelencia Severo Ochoa (CEX2021-001165-S), and co-financed by the European Regional Development Fund (ERDF).

---

**Figure S1.**
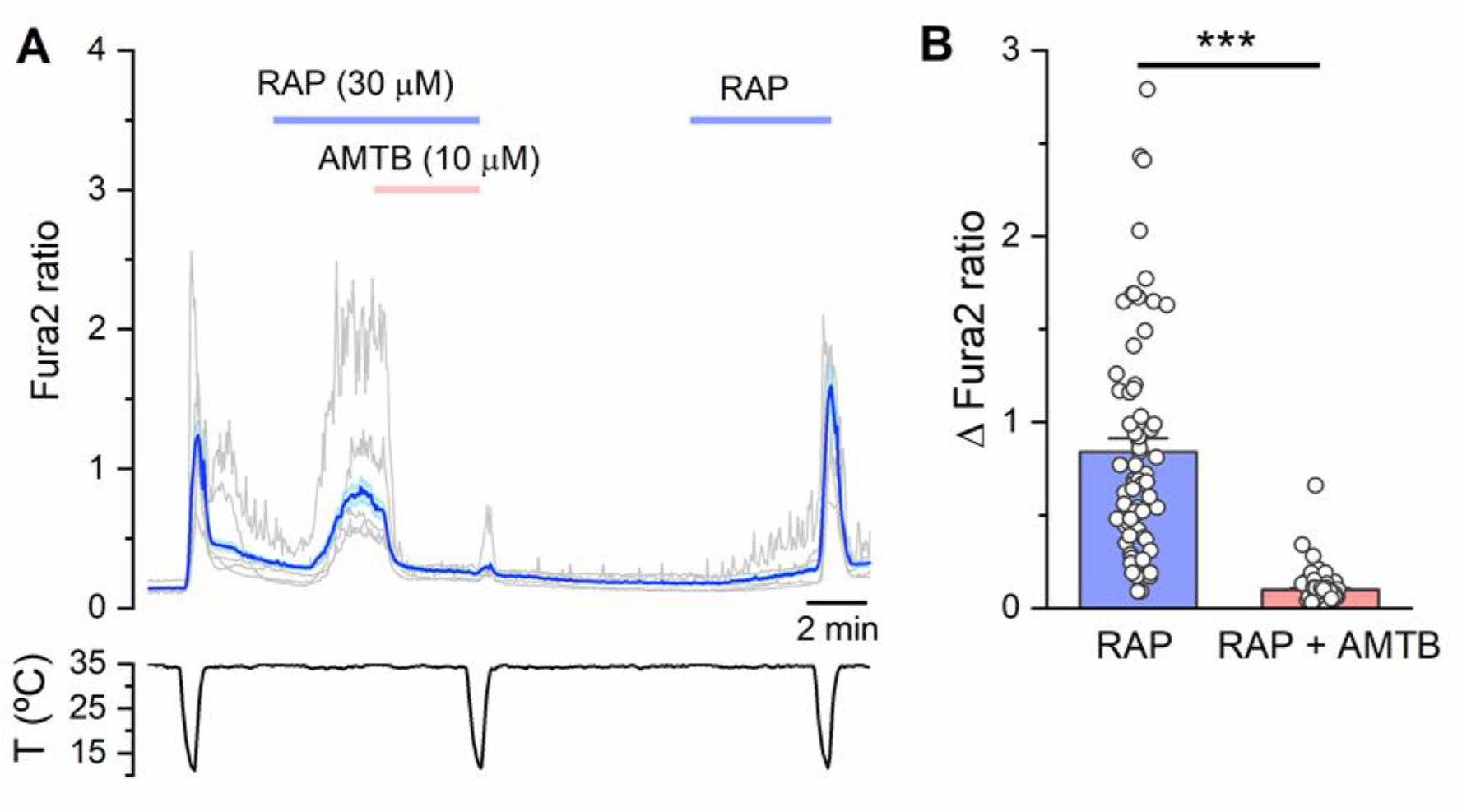
Rapamycin responses are mediated by TRPM8. **A.** Time course of intracellular calcium responses in HEK293 cells transfected with mouse TRPM8. The application of AMTB blocks responses to cold and to RAP. The blue trace represents the average ± SEM of the response. The grey traces represent a sample of single cell responses. **B**. Summary histogram of the effect of AMTB on rapamycin-evoked individual and average response. Paired Student t-test (n = 69, 2 experiments).

**Figure S2.**
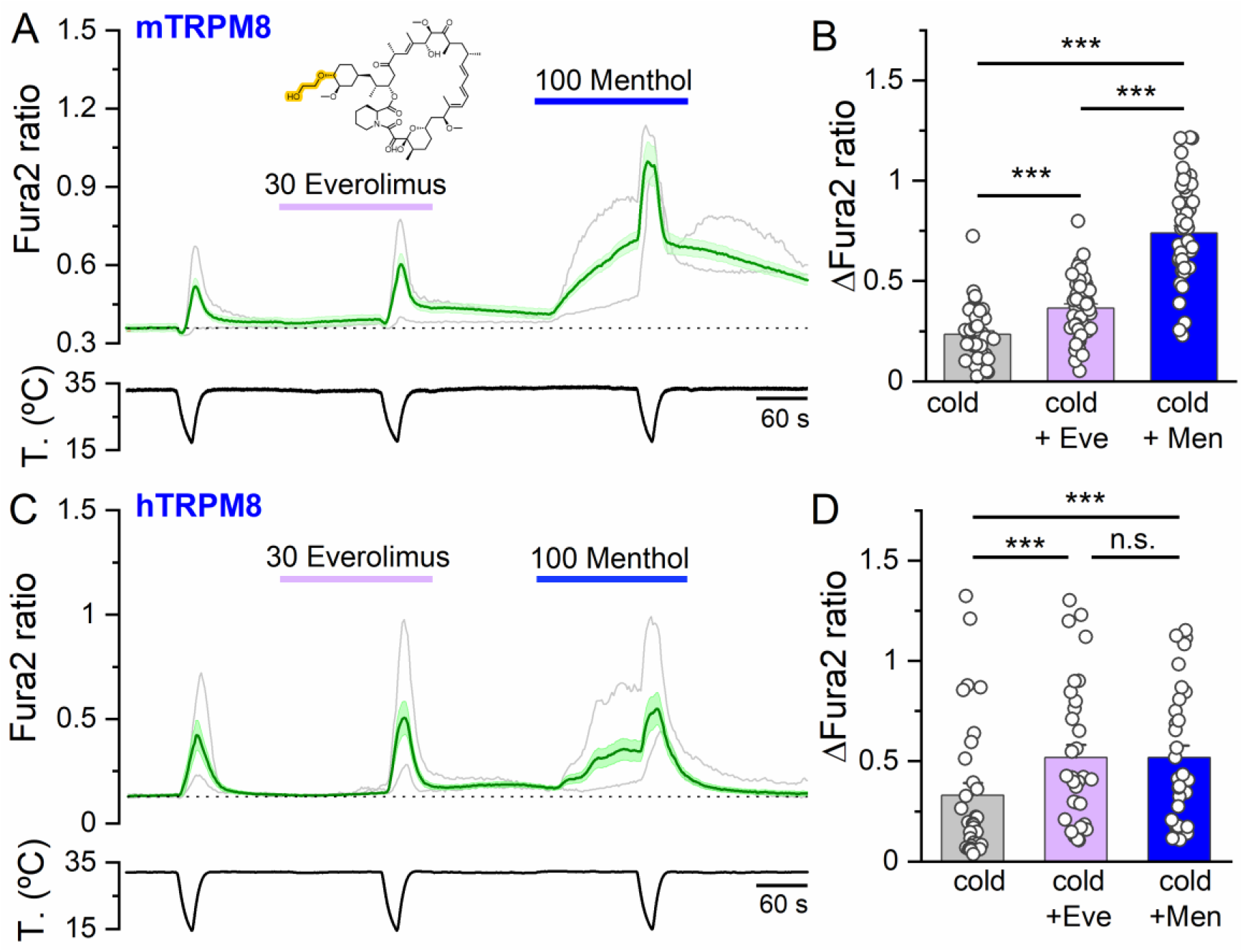
Everolimus is a weak TRPM8 agonist. **A.** Time course of intracellular calcium responses in HEK293 cells transfected with mouse TRPM8. Everolimus (30 µM), produced an insignificant response compared to menthol (100 µM), but clearly potentiated cold-evoked responses. The green trace represents the average ± SEM response on 16 cells in the field. The grey traces represent examples of individual cells. **B**. Summary histogram of the cold-evoked individual and mean responses in control solution, everolimus and menthol. Average ± SEM of 48 cells in 3 experiments. **C**. The same experiment in HEK293 cells transfected with human TRPM8. B. Summary histogram of the cold-evoked responses. **D.** Summary histogram of the cold-evoked responses in control solution, everolimus and menthol. Average ± SEM of 32 cells in 2 experiments. Statistical significance calculated by a one-way ANOVA followed by Bonferroni post-test.

**Figure S3.**
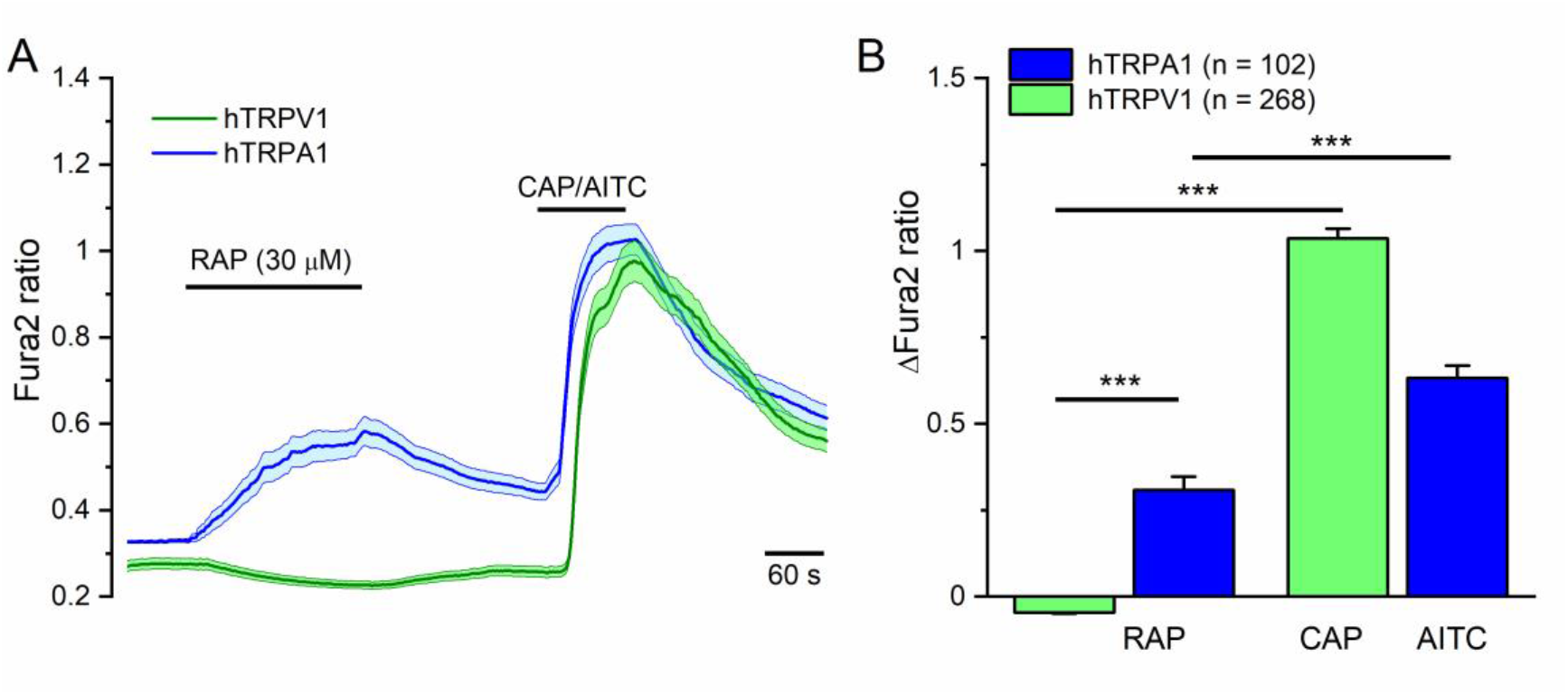
Effects of rapamycin on other human thermoTRP channels. **A.** Time course of intracellular calcium responses in HEK293 cells transfected with human TRPV1 or human TRPA1, stimulated with RAP (30 μM) or its canonical agonist (Capsaicin 100 nM or AITC 100 µM). The green and blue traces represent the average ± SEM response for TRPV1- and TRPA1-transfected cells respectively. RAP is a weak TRPA1 agonist and TRPV1 is not activated by RAP. **B.** Summary histogram of the responses to RAP and capsaicin/AITC. Average ± SEM of 268 hTRPV1 (3 experiments) and 102 hTRPA1 cells (2 experiments) respectively. Statistical significance calculated by a one-way ANOVA followed by Bonferroni post-test.

**Figure S4.**
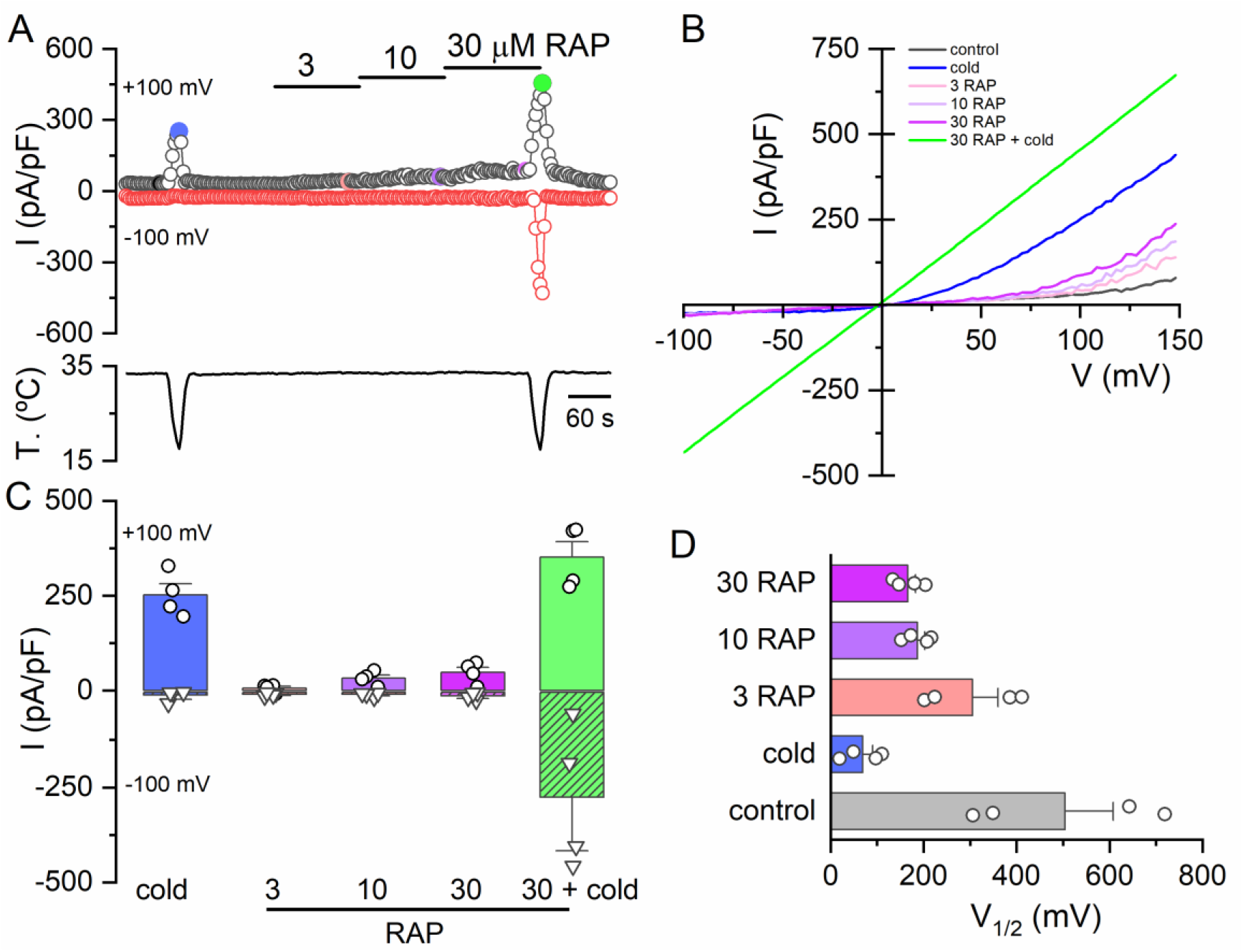
Rapamycin activates TRPM8 at low concentrations. ***A,*** Representative time course of whole-cell current at −100 and +100 mV in a HEK293 cell transiently transfected with mTRPM8 during the sequential application of increasing concentrations of rapamycin. The bottom trace shows the simultaneous recording of the bath temperature during the experiment. ***B,*** Current-voltage (I-V) relationship obtained by a 400 ms voltage ramp from −100 to +150 mV during the experiment shown in A. The color of individual traces matches the color at each particular time point in A. ***C,*** Bar histogram of individual and mean ± SEM current density values at −100 and +100 mV evoked by the different stimuli shown in A, with the same color code (n = 4 cells, 1 experiment). ***D,*** Individual and mean ± SEM V_1/2_ values calculated from fitting individual I-V curves to a linearized Boltzmann equation.

**Figure S5.**
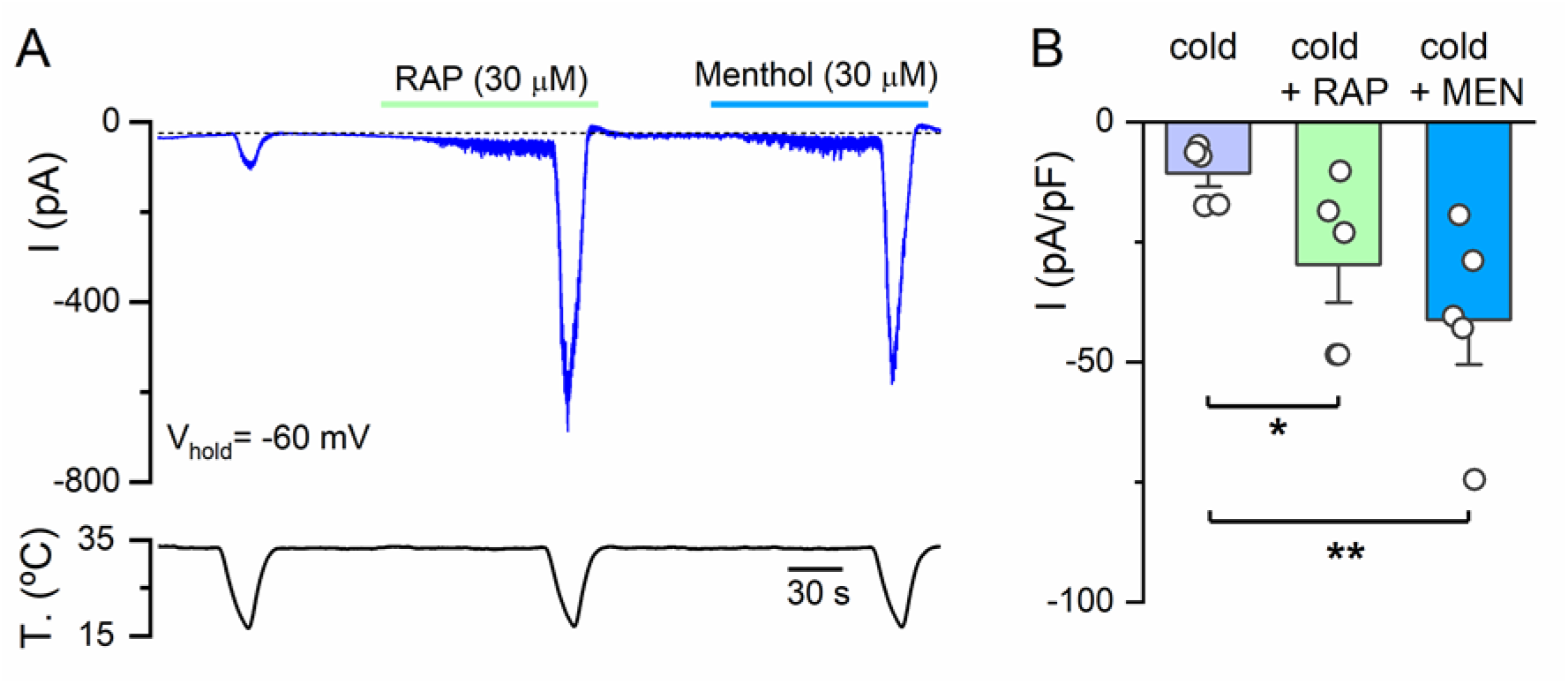
Rapamycin and menthol activate cold-sensitive DRG neurons. **A,** Whole-cell recording in the voltage-clamp configuration (V_hold_= −60 mV) of a TRPM8-expressing, cold-sensitive DRG neuron identified by EYFP expression. Bottom trace is the recording of bath temperature during the experiment. RAP and menthol activate similar small inward currents and strongly potentiate the response to cold. ***B,*** Bar histogram of individual and mean peak inward current density evoked by cold, RAP plus cold and menthol plus cold (n = 5, 1 experiment). The statistical analysis consisted of a one-way ANOVA for repeated measures followed by Tukey post-hoc test.

## References

Almaraz, L., Manenschijn, J. A., de la Pena, E., & Viana, F. (2014). Trpm8. Handb Exp Pharmacol, 222, 547–579. 10.1007/978-3-642-54215-2_22

Arcas, J. M., Gonzalez, A., Gers-Barlag, K., Gonzalez-Gonzalez, O., Bech, F., Demirkhanyan, L., … Viana, F. (2019). The Immunosuppressant Macrolide Tacrolimus Activates Cold-Sensing TRPM8 Channels. J Neurosci, 39(6), 949–969. 10.1523/JNEUROSCI.1726-18.2018

Asuthkar, S., Demirkhanyan, L., Sun, X., Elustondo, P. A., Krishnan, V., Baskaran, P., … Zakharian, E. (2015). The TRPM8 protein is a testosterone receptor: II. Functional evidence for an ionotropic effect of testosterone on TRPM8. J Biol Chem, 290(5), 2670–2688. 10.1074/jbc.M114.610873

Bandell, M., Dubin, A. E., Petrus, M. J., Orth, A., Mathur, J., Hwang, S. W., & Patapoutian, A. (2006). High-throughput random mutagenesis screen reveals TRPM8 residues specifically required for activation by menthol. Nat Neurosci, 9(4), 493–500. 10.1038/nn1665

Benjamin, D., Colombi, M., Moroni, C., & Hall, M. N. (2011). Rapamycin passes the torch: a new generation of mTOR inhibitors. Nat Rev Drug Discov, 10(11), 868–880. 10.1038/nrd3531

Bidaux, G., Borowiec, A. S., Dubois, C., Delcourt, P., Schulz, C., Vanden Abeele, F., … Prevarskaya, N. (2016). Targeting of short TRPM8 isoforms induces 4TM-TRPM8-dependent apoptosis in prostate cancer cells. Oncotarget, 7(20), 29063–29080. 10.18632/oncotarget.8666

Bidaux, G., Borowiec, A. S., Gordienko, D., Beck, B., Shapovalov, G. G., Lemonnier, L., … Prevarskaya, N. (2015). Epidermal TRPM8 channel isoform controls the balance between keratinocyte proliferation and differentiation in a cold-dependent manner. Proc Natl Acad Sci U S A, 112(26), E3345–3354. 10.1073/pnas.1423357112

Bitto, A., Ito, T. K., Pineda, V. V., LeTexier, N. J., Huang, H. Z., Sutlief, E., … Kaeberlein, M. (2016). Transient rapamycin treatment can increase lifespan and healthspan in middle-aged mice. Elife, 5. 10.7554/eLife.16351

Brauchi, S., Orio, P., & Latorre, R. (2004). Clues to understanding cold sensation: thermodynamics and electrophysiological analysis of the cold receptor TRPM8. Proc Natl Acad Sci U S A, 101(43), 15494–15499. 10.1073/pnas.0406773101

Brillantes, A. B., Ondrias, K., Scott, A., Kobrinsky, E., Ondriasova, E., Moschella, M. C., … Marks, A. R. (1994). Stabilization of calcium release channel (ryanodine receptor) function by FK506-binding protein. Cell, 77(4), 513–523. https://www.ncbi.nlm.nih.gov/pubmed/7514503

Choi, J., Chen, J., Schreiber, S. L., & Clardy, J. (1996). Structure of the FKBP12-rapamycin complex interacting with the binding domain of human FRAP. Science, 273(5272), 239–242. 10.1126/science.273.5272.239

Conti, B. (2008). Considerations on temperature, longevity and aging. Cell Mol Life Sci, 65(11), 1626–1630. 10.1007/s00018-008-7536-1

Conti, B., Sanchez-Alavez, M., Winsky-Sommerer, R., Morale, M. C., Lucero, J., Brownell, S., … Bartfai, T. (2006). Transgenic mice with a reduced core body temperature have an increased life span. Science, 314(5800), 825–828. 10.1126/science.1132191

Denda, M., Tsutsumi, M., & Denda, S. (2010). Topical application of TRPM8 agonists accelerates skin permeability barrier recovery and reduces epidermal proliferation induced by barrier insult: role of cold-sensitive TRP receptors in epidermal permeability barrier homoeostasis. Exp Dermatol, 19(9), 791–795. 10.1111/j.1600-0625.2010.01154.x

Dhaka, A., Earley, T. J., Watson, J., & Patapoutian, A. (2008). Visualizing cold spots: TRPM8-expressing sensory neurons and their projections. J Neurosci, 28(3), 566–575. 10.1523/JNEUROSCI.3976-07.2008

Dhaka, A., Murray, A. N., Mathur, J., Earley, T. J., Petrus, M. J., & Patapoutian, A. (2007). TRPM8 is required for cold sensation in mice. Neuron, 54(3), 371–378. 10.1016/j.neuron.2007.02.024

Foster, R. S., Bint, L. J., & Halbert, A. R. (2012). Topical 0.1% rapamycin for angiofibromas in paediatric patients with tuberous sclerosis: a pilot study of four patients. Australas J Dermatol, 53(1), 52–56. 10.1111/j.1440-0960.2011.00837.x

Hammond, G. R., Fischer, M. J., Anderson, K. E., Holdich, J., Koteci, A., Balla, T., & Irvine, R. F. (2012). PI4P and PI(4,5)P2 are essential but independent lipid determinants of membrane identity. Science, 337(6095), 727–730. 10.1126/science.1222483

Harrison, D. E., Strong, R., Sharp, Z. D., Nelson, J. F., Astle, C. M., Flurkey, K., … Miller, R. A. (2009). Rapamycin fed late in life extends lifespan in genetically heterogeneous mice. Nature, 460(7253), 392–395. 10.1038/nature08221

Heitman, J., Movva, N. R., & Hall, M. N. (1991). Targets for cell cycle arrest by the immunosuppressant rapamycin in yeast. Science, 253(5022), 905–909. 10.1126/science.1715094

Houghton, P. J. (2010). Everolimus. Clin Cancer Res, 16(5), 1368–1372. 10.1158/1078-0432.CCR-09-1314

Izquierdo, C., Martin-Martinez, M., Gomez-Monterrey, I., & Gonzalez-Muniz, R. (2021). TRPM8 Channels: Advances in Structural Studies and Pharmacological Modulation. Int J Mol Sci, 22(16). 10.3390/ijms22168502

Janssens, A., Gees, M., Toth, B. I., Ghosh, D., Mulier, M., Vennekens, R., … Voets, T. (2016). Definition of two agonist types at the mammalian cold-activated channel TRPM8. Elife, 5. 10.7554/eLife.17240

Johnson, S. C., Rabinovitch, P. S., & Kaeberlein, M. (2013). mTOR is a key modulator of ageing and age-related disease. Nature, 493(7432), 338–345. 10.1038/nature11861

Julius, D. (2005). From peppers to peppermints: natural products as probes of the pain pathway. Harvey Lect, 101, 89–115. https://www.ncbi.nlm.nih.gov/pubmed/18030977

Kennedy, B. K., & Lamming, D. W. (2016). The Mechanistic Target of Rapamycin: The Grand ConducTOR of Metabolism and Aging. Cell Metab, 23(6), 990–1003. 10.1016/j.cmet.2016.05.009

Kita, T., Uchida, K., Kato, K., Suzuki, Y., Tominaga, M., & Yamazaki, J. (2019). FK506 (tacrolimus) causes pain sensation through the activation of transient receptor potential ankyrin 1 (TRPA1) channels. J Physiol Sci, 69(2), 305–316. 10.1007/s12576-018-0647-z

Knowlton, W. M., Palkar, R., Lippoldt, E. K., McCoy, D. D., Baluch, F., Chen, J., & McKemy, D. D. (2013). A sensory-labeled line for cold: TRPM8-expressing sensory neurons define the cellular basis for cold, cold pain, and cooling-mediated analgesia. J Neurosci, 33(7), 2837–2848. 10.1523/JNEUROSCI.1943-12.2013

Kunz, J., & Hall, M. N. (1993). Cyclosporin A, FK506 and rapamycin: more than just immunosuppression. Trends Biochem Sci, 18(9), 334–338. https://www.ncbi.nlm.nih.gov/pubmed/7694398

Lashinger, E. S., Steiginga, M. S., Hieble, J. P., Leon, L. A., Gardner, S. D., Nagilla, R., … Su, X. (2008). AMTB, a TRPM8 channel blocker: evidence in rats for activity in overactive bladder and painful bladder syndrome. Am J Physiol Renal Physiol, 295(3), F803–810. 10.1152/ajprenal.90269.2008

Li, J., Kim, S. G., & Blenis, J. (2014). Rapamycin: one drug, many effects. Cell Metab, 19(3), 373–379. 10.1016/j.cmet.2014.01.001

Liu, B., Fan, L., Balakrishna, S., Sui, A., Morris, J. B., & Jordt, S. E. (2013). TRPM8 is the principal mediator of menthol-induced analgesia of acute and inflammatory pain. Pain, 154(10), 2169–2177. 10.1016/j.pain.2013.06.043

Loeb, J., & Northrop, J. H. (1916). Is There a Temperature Coefficient for the Duration of Life? Proc Natl Acad Sci U S A, 2(8), 456–457. 10.1073/pnas.2.8.456

Lombardi, A., Trimarco, B., Iaccarino, G., & Santulli, G. (2017). Impaired mitochondrial calcium uptake caused by tacrolimus underlies beta-cell failure. Cell Commun Signal, 15(1), 47. 10.1186/s12964-017-0203-0

Madrid, R., de la Pena, E., Donovan-Rodriguez, T., Belmonte, C., & Viana, F. (2009). Variable threshold of trigeminal cold-thermosensitive neurons is determined by a balance between TRPM8 and Kv1 potassium channels. J Neurosci, 29(10), 3120–3131. 10.1523/JNEUROSCI.4778-08.2009

Malkia, A., Pertusa, M., Fernandez-Ballester, G., Ferrer-Montiel, A., & Viana, F. (2009). Differential role of the menthol-binding residue Y745 in the antagonism of thermally gated TRPM8 channels. Mol Pain, 5, 62. 10.1186/1744-8069-5-62

Mangal, S., Zielich, J., Lambie, E., & Zanin, E. (2018). Rapamycin-induced protein dimerization as a tool for C. elegans research. MicroPubl Biol, 2018. 10.17912/W2BH3H

Martinez-Lopez, P., Trevino, C. L., de la Vega-Beltran, J. L., De Blas, G., Monroy, E., Beltran, C., … Darszon, A. (2011). TRPM8 in mouse sperm detects temperature changes and may influence the acrosome reaction. J Cell Physiol, 226(6), 1620–1631. 10.1002/jcp.22493

McKemy, D. D., Neuhausser, W. M., & Julius, D. (2002). Identification of a cold receptor reveals a general role for TRP channels in thermosensation. Nature, 416(6876), 52–58. 10.1038/nature719

Meotti, F. C., Lemos de Andrade, E., & Calixto, J. B. (2014). TRP modulation by natural compounds. Handb Exp Pharmacol, 223, 1177–1238. 10.1007/978-3-319-05161-1_19

Moran, M. M., & Szallasi, A. (2017). Targeting nociceptive transient receptor potential channels to treat chronic pain: current state of the field. Br J Pharmacol. 10.1111/bph.14044

Morenilla-Palao, C., Luis, E., Fernandez-Pena, C., Quintero, E., Weaver, J. L., Bayliss, D. A., & Viana, F. (2014). Ion channel profile of TRPM8 cold receptors reveals a role of TASK-3 potassium channels in thermosensation. Cell Rep, 8(5), 1571–1582. 10.1016/j.celrep.2014.08.003

Mutizwa, M. M., Berk, D. R., & Anadkat, M. J. (2011). Treatment of facial angiofibromas with topical application of oral rapamycin solution (1mgmL(−1)) in two patients with tuberous sclerosis. Br J Dermatol, 165(4), 922–923. 10.1111/j.1365-2133.2011.10476.x

Nilius, B., & Appendino, G. (2011). Tasty and healthy TR(i)Ps. The human quest for culinary pungency. EMBO Rep, 12(11), 1094–1101. 10.1038/embor.2011.200

Olivares, E., Salgado, S., Maidana, J. P., Herrera, G., Campos, M., Madrid, R., & Orio, P. (2015). TRPM8-Dependent Dynamic Response in a Mathematical Model of Cold Thermoreceptor. PLoS One, 10(10), e0139314. 10.1371/journal.pone.0139314

Palkar, R., Ongun, S., Catich, E., Li, N., Borad, N., Sarkisian, A., & McKemy, D. D. (2018). Cooling Relief of Acute and Chronic Itch Requires TRPM8 Channels and Neurons. J Invest Dermatol, 138(6), 1391–1399. 10.1016/j.jid.2017.12.025

Parra, A., Madrid, R., Echevarria, D., del Olmo, S., Morenilla-Palao, C., Acosta, M. C., … Belmonte, C. (2010). Ocular surface wetness is regulated by TRPM8-dependent cold thermoreceptors of the cornea. Nat Med, 16(12), 1396–1399. 10.1038/nm.2264

Peier, A. M., Moqrich, A., Hergarden, A. C., Reeve, A. J., Andersson, D. A., Story, G. M., … Patapoutian, A. (2002). A TRP channel that senses cold stimuli and menthol. Cell, 108(5), 705–715. 10.1016/s0092-8674(02)00652-9

Proudfoot, C. J., Garry, E. M., Cottrell, D. F., Rosie, R., Anderson, H., Robertson, D. C., … Mitchell, R. (2006). Analgesia mediated by the TRPM8 cold receptor in chronic neuropathic pain. Curr Biol, 16(16), 1591–1605. 10.1016/j.cub.2006.07.061

Quallo, T., Vastani, N., Horridge, E., Gentry, C., Parra, A., Moss, S., … Bevan, S. (2015). TRPM8 is a neuronal osmosensor that regulates eye blinking in mice. Nat Commun, 6, 7150. 10.1038/ncomms8150

Reid, G., Amuzescu, B., Zech, E., & Flonta, M. L. (2001). A system for applying rapid warming or cooling stimuli to cells during patch clamp recording or ion imaging. J Neurosci Methods, 111(1), 1–8. https://www.ncbi.nlm.nih.gov/pubmed/11574114

Reid, G., Babes, A., & Pluteanu, F. (2002). A cold- and menthol-activated current in rat dorsal root ganglion neurones: properties and role in cold transduction. J Physiol, 545(2), 595–614. 10.1113/jphysiol.2002.024331

Reimundez, A., Fernandez-Pena, C., Garcia, G., Fernandez, R., Ordas, P., Gallego, R., … Senaris, R. (2018). Deletion of the Cold Thermoreceptor TRPM8 Increases Heat Loss and Food Intake Leading to Reduced Body Temperature and Obesity in Mice. J Neurosci, 38(15), 3643–3656. 10.1523/JNEUROSCI.3002-17.2018

Rivera, B., Moreno, C., Lavanderos, B., Hwang, J. Y., Fernandez-Trillo, J., Park, K. S., … Pertusa, M. (2021). Constitutive Phosphorylation as a Key Regulator of TRPM8 Channel Function. J Neurosci, 41(41), 8475–8493. 10.1523/JNEUROSCI.0345-21.2021

Rovira, J., Diekmann, F., Ramirez-Bajo, M. J., Banon-Maneus, E., Moya-Rull, D., & Campistol, J. M. (2012). Sirolimus-associated testicular toxicity: detrimental but reversible. Transplantation, 93(9), 874–879. 10.1097/TP.0b013e31824bf1f0

Roza, C., Belmonte, C., & Viana, F. (2006). Cold sensitivity in axotomized fibers of experimental neuromas in mice. Pain, 120(1-2), 24–35. 10.1016/j.pain.2005.10.006

Sarria, I., Ling, J., Zhu, M. X., & Gu, J. G. (2011). TRPM8 acute desensitization is mediated by calmodulin and requires PIP(2): distinction from tachyphylaxis. J Neurophysiol, 106(6), 3056–3066. 10.1152/jn.00544.2011

Saxton, R. A., & Sabatini, D. M. (2017). mTOR Signaling in Growth, Metabolism, and Disease. Cell, 168(6), 960–976. 10.1016/j.cell.2017.02.004

Seto, B. (2012). Rapamycin and mTOR: a serendipitous discovery and implications for breast cancer. Clin Transl Med, 1(1), 29. 10.1186/2001-1326-1-29

Spatola, R., Nadelstein, B., Berdoulay, A., & English, R. V. (2018). The effects of topical aqueous sirolimus on tear production in normal dogs and dogs with refractory dry eye. Vet Ophthalmol, 21(3), 255–263. 10.1111/vop.12503

Takashima, Y., Daniels, R. L., Knowlton, W., Teng, J., Liman, E. R., & McKemy, D. D. (2007). Diversity in the neural circuitry of cold sensing revealed by genetic axonal labeling of transient receptor potential melastatin 8 neurons. J Neurosci, 27(51), 14147–14157. 10.1523/JNEUROSCI.4578-07.2007

Toro, C. A., Eger, S., Veliz, L., Sotelo-Hitschfeld, P., Cabezas, D., Castro, M. A., … Brauchi, S. (2015). Agonist-dependent modulation of cell surface expression of the cold receptor TRPM8. J Neurosci, 35(2), 571–582. 10.1523/JNEUROSCI.3820-13.2015

Tsavaler, L., Shapero, M. H., Morkowski, S., & Laus, R. (2001). Trp-p8, a novel prostate-specific gene, is up-regulated in prostate cancer and other malignancies and shares high homology with transient receptor potential calcium channel proteins. Cancer Res, 61(9), 3760–3769. https://www.ncbi.nlm.nih.gov/pubmed/11325849

van Unen, J., Rashidfarrokhi, A., Hoogendoorn, E., Postma, M., Gadella, T. W., Jr., & Goedhart, J. (2016). Quantitative Single-Cell Analysis of Signaling Pathways Activated Immediately Downstream of Histamine Receptor Subtypes. Mol Pharmacol, 90(3), 162–176. 10.1124/mol.116.104505

Varnai, P., Thyagarajan, B., Rohacs, T., & Balla, T. (2006). Rapidly inducible changes in phosphatidylinositol 4,5-bisphosphate levels influence multiple regulatory functions of the lipid in intact living cells. J Cell Biol, 175(3), 377–382. 10.1083/jcb.200607116

Vezina, C., Kudelski, A., & Sehgal, S. N. (1975). Rapamycin (AY-22,989), a new antifungal antibiotic. I. Taxonomy of the producing streptomycete and isolation of the active principle. J Antibiot (Tokyo), 28(10), 721–726. 10.7164/antibiotics.28.721

Voets, T., Droogmans, G., Wissenbach, U., Janssens, A., Flockerzi, V., & Nilius, B. (2004). The principle of temperature-dependent gating in cold- and heat-sensitive TRP channels. Nature, 430(7001), 748–754. 10.1038/nature02732

Wilkinson, J. E., Burmeister, L., Brooks, S. V., Chan, C. C., Friedline, S., Harrison, D. E., … Miller, R. A. (2012). Rapamycin slows aging in mice. Aging Cell, 11(4), 675–682. 10.1111/j.1474-9726.2012.00832.x

Winter, Z., Gruschwitz, P., Eger, S., Touska, F., & Zimmermann, K. (2017). Cold Temperature Encoding by Cutaneous TRPA1 and TRPM8-Carrying Fibers in the Mouse. Front Mol Neurosci, 10, 209. 10.3389/fnmol.2017.00209

Wirta, D. L., Senchyna, M., Lewis, A. E., Evans, D. G., McLaurin, E. B., Ousler, G. W., & Hollander, D. A. (2022). A randomized, vehicle-controlled, Phase 2b study of two concentrations of the TRPM8 receptor agonist AR-15512 in the treatment of dry eye disease (COMET-1). Ocul Surf, 26, 166–173. 10.1016/j.jtos.2022.08.003

Xiao, R., Zhang, B., Dong, Y., Gong, J., Xu, T., Liu, J., & Xu, X. Z. (2013). A genetic program promotes C. elegans longevity at cold temperatures via a thermosensitive TRP channel. Cell, 152(4), 806–817. 10.1016/j.cell.2013.01.020

Yamamura, H., Ugawa, S., Ueda, T., Morita, A., & Shimada, S. (2008). TRPM8 activation suppresses cellular viability in human melanoma. Am J Physiol Cell Physiol, 295(2), C296–301. 10.1152/ajpcell.00499.2007

Yang, J. M., Li, F., Liu, Q., Ruedi, M., Wei, E. T., Lentsman, M., … Yoon, K. C. (2017). A novel TRPM8 agonist relieves dry eye discomfort. BMC Ophthalmol, 17(1), 101. 10.1186/s12886-017-0495-2

Zakharian, E., Cao, C., & Rohacs, T. (2010). Gating of transient receptor potential melastatin 8 (TRPM8) channels activated by cold and chemical agonists in planar lipid bilayers. J Neurosci, 30(37), 12526–12534. 10.1523/JNEUROSCI.3189-10.2010

Zakharian, E., Thyagarajan, B., French, R. J., Pavlov, E., & Rohacs, T. (2009). Inorganic polyphosphate modulates TRPM8 channels. PLoS One, 4(4), e5404. 10.1371/journal.pone.0005404

Zhang, L., & Barritt, G. J. (2006). TRPM8 in prostate cancer cells: a potential diagnostic and prognostic marker with a secretory function? Endocr Relat Cancer, 13(1), 27–38. 10.1677/erc.1.01093

Zhang, X. (2019). Direct Galpha(q) Gating Is the Sole Mechanism for TRPM8 Inhibition Caused by Bradykinin Receptor Activation. Cell Rep, 27(12), 3672–3683 e3674. 10.1016/j.celrep.2019.05.080

Zhang, X., Chen, W., Gao, Q., Yang, J., Yan, X., Zhao, H., … Xu, H. (2019). Rapamycin directly activates lysosomal mucolipin TRP channels independent of mTOR. PLoS Biol, 17(5), e3000252. 10.1371/journal.pbio.3000252

Zimmermann, K., Hein, A., Hager, U., Kaczmarek, J. S., Turnquist, B. P., Clapham, D. E., & Reeh, P. W. (2009). Phenotyping sensory nerve endings in vitro in the mouse. Nat Protoc, 4(2), 174–196. 10.1038/nprot.2008.223

Zou, Z., Tao, T., Li, H., & Zhu, X. (2020). mTOR signaling pathway and mTOR inhibitors in cancer: progress and challenges. Cell Biosci, 10, 31. 10.1186/s13578-020-00396-1

